# A rhizarian genome reveals an osmotrophic route to extracellular digestion in eukaryotes

**DOI:** 10.64898/2026.07.16.738196

**Authors:** Hüsna Öztoprak, Julia Graf, Robin Jacobs, Fiona Kaiser, Kenneth Dumack

## Abstract

Rhizaria, one of the most species-rich and ecologically important eukaryotic supergroups, accommodates a newly discovered osmotrophic species, yet high-quality genomic resources remain scarce, limiting our understanding of their metabolic diversity and ecological functions. Here, we present the genome of *Saccharomycomorpha psychra*, the first rhizarian telomere-anchored assembly, to be explored as a model and reference for rhizarian ecology and evolution. The 62 Mb assembly, of which half comprises 22 telomere-to-telomere scaffolds, shows a BUSCO completeness of 94.2%, encodes 17,680 genes, and provides the genomic foundation for investigating rhizarian ecology and evolution. The genome of osmotrophic *Saccharomycomorpha psychra* reveals a functionally integrated secretome of 1,015 proteins dominated by carbohydrate-active enzymes (CAZymes), proteases, lipases, and oxidoreductases. Taken together with 303 predicted high-confidence membrane transporters of a total of 680, skewed toward H⁺-coupled secondary carriers, these features constitute the genomic signature of an extracellular digestive strategy convergent with saprotrophic fungi. Phenotypic MicroArray**^TM^** assays confirmed active utilization of 17 carbon sources, including all six C5 pentose sugars tested, consistent with the predicted arabinose and ribokinase pathways among the most highly expressed metabolic genes in the transcriptome. These findings demonstrate that osmotrophic saprotrophy has evolved independently in Rhizaria with enzymatic solutions that closely parallel those of evolutionary distant fungal decomposers, highlighting the power of ecological context over phylogenetic heritage in shaping extracellular metabolic architecture.

## Introduction

The vast diversity of fungi and bacteria are osmotrophic, i.e. degrade environmental compounds via secreted enzymes. In contrast, most heterotrophic microbial eukaryotes phagocytose prey (López-García & Moreira, 2026; Ocaña-Pallarès et al., 2026). Rhizaria are a major eukaryotic supergroup comprising diverse protists with functional versatility driving global biogeochemistry. In the oceans, Foraminifera, Radiolaria, and Phaeodaria, contribute to overall environmental chemistry through predation and through biomineralization to silica cycles and fossil formation. In soil, Rhizaria represent a majority of protists, where Sarcomonadea (Rhizaria) alone represents approximately 10% to 50% of all recovered protistan reads (Oliverio et al., 2020). Their high abundance, short generation times, phagotrophic predatory behavior and small size suggests they play a key role in the soil food web by channeling energy to higher trophic levels (Azam et al., 1983; Clarholm, 1985; de Ruiter et al., 1995). Furthermore, rhizarian parasite *Plasmodiophora brassicae* causes about 15% losses in cabbage yields worldwide (Cavalier-Smith et al., 2018; Dumack et al., 2019; Nakamura & Suzuki, 2015; Neuhauser et al., 2010; Nowack, 2014).

Despite their significance, Rhizaria is the only major supergroup within eukaryotic diversity which is abundant, ubiquitous, largely diverse, but had lacked any high-quality, chromosome-scale genome and thus functional, metabolic understanding. In bacteriology, genomes provided valuable information on their functions, including an improved understanding of processes such as nitrogen cycling, fermentation, and the spread of antimicrobial resistance (Collineau et al., 2019; Herold et al., 2020; Imachi et al., 2020; Zehr & Capone, 2020). In eukaryotic microbiology, available genomic datasets are both scarce and biased towards certain functional groups. The majority of the <600 sequenced protistan genomes stem from respective parasitic or algal groups (Schoenle et al., 2025; Sibbald & Archibald, 2017).

Here, we provide the first genomic and metabolic exploration of a free-living and heterotrophic rhizarian species, *Saccharomycomorpha psychra*. *S. psychra* was only recently described (Feng et al., 2021). In contrast to its relatives, it lacks any evidence of phagocytosis and instead facilitates high-density, axenic culturing on agar, supporting high-quality genome sequencing and experimental verification of metabolic capabilities. Based on this, we explore the rhizarian metabolic versatility, secretome, and membrane transport system. As *S. psychra* was isolated from Antarctic lichen, a structurally and chemically heterogeneous biopolymer-rich habitat shared with fungi, bacteria, and other microbial eukaryotes, we further ask whether its secreted enzyme repertoire reflects ecological convergence with other lichen-associated decomposers, rather than shared ancestry with its phagotrophic rhizarian relatives.

## Results

### A telomere-anchored rhizarian genome provides a foundation for metabolic reconstruction

We assembled a rhizarian reference-quality genome of *Saccharomycomorpha psychra* (Fig. S1), isolated from Antarctic lichen. Pre-assembly *k*-mer analysis of PacBio HiFi reads indicated a diploid genome with heterozygosity of 9.2% (Fig. S2), consistent with the complexity observed during assembly. The final assembly spans 61.7 Mb and comprises 78 scaffolds with an N_50_ of 1.6 Mb, including 22 chromosome-level scaffolds, defined by telomeric repeats (‘AACCCT’) at both termini and, representing 54.8% of the assembly length (Fig. 1). An additional 26 scaffolds contain telomeric repeats at one end, increasing nearly chromosome-scale sequence representation to 51.1 Mb (82.9% of the assembly). Genome completeness was high, with BUSCO scores of 94.2% against the Alveolata database (90.7% single-copy orthologs, 3.5% duplicated orthologs) and 81.2% against the broader Eukaryota database (79.2% single-copy orthologs, 2.0% duplicated orthologs); note a Rhizaria database does not exist due to the yet existing lack of sufficient reference data (Fig. S3, S4). Structural annotation predicted 17,680 protein-coding genes, providing a gene-rich framework for functional reconstruction. The genome contains an intron density of 0.73 introns per gene, repeat content of 34.15% and transposable elements (TEs) accounted for 19.07% of the assembly, with LTR retrotransposons (RLC/Copia, RLG/Gypsy) as the dominant classes during a burst event (Table S1, Fig. S5, S6). The genome is markedly gene-dense, with TE density negatively correlated with both exon and gene density (Fig.1, S5). Together, these assembly and annotation metrics establish *S. psychra* as the highest-quality genomic reference for investigating the molecular basis of osmotrophy in Rhizaria (Table S2).

**Figure 1:**
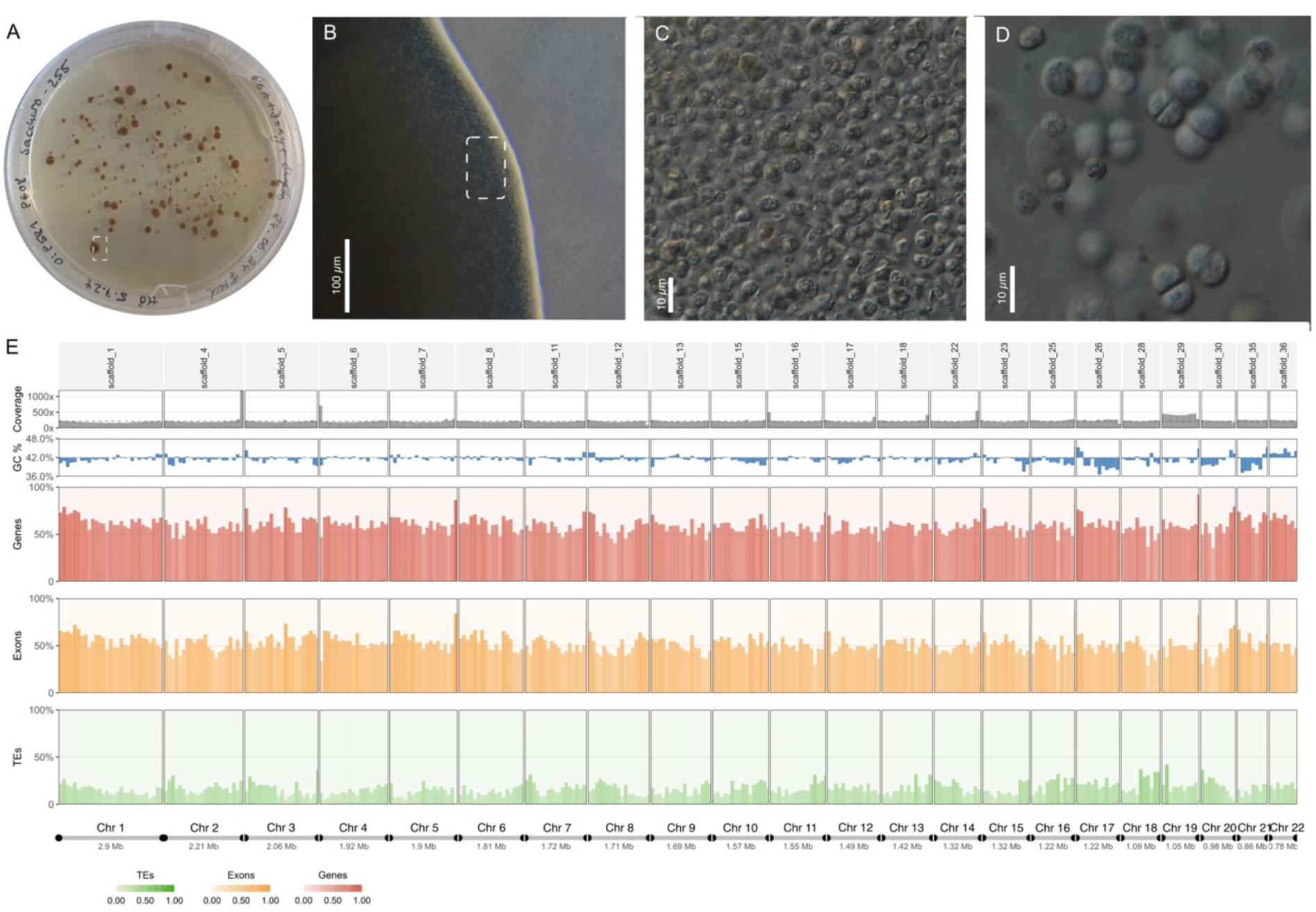
Morphology and telomere-anchored genome assembly architecture of *Saccharomycomorpha psychra*. **A-D:** Morphology of *S. psychra* across increasing magnification. Dashed boxes in (A) and (B) indicate the region shown at higher magnification in the subsequent panel. (**A)** colony morphology on solid medium. **(B)** Light micrograph of densely aggregated cells. **(C, D)**: Higher magnification view of the cellular assemblage and individual cells within a single-layer region of assemblage. **(E)** Representation of the genome assembly with twenty-two telomere-to-telomere scaffolds shown in descending order of length, representing 54.8% of the assembled genome. Tracks indicate read coverage (grey), GC content (blue), gene density (red), exon density (orange) and transposable-element (TE) density (green), calculated in 100-kb windows. Color intensity for gene, exon and TE tracks correspond to the proportion of bases occupied within each window. Telomeric regions are indicated as black markers.

### Central metabolism supports aerobic processing of imported dissolved carbon

Functional annotation recovered 9,968 genes with KEGG Orthology assignments, of which 96.4% were transcriptionally detected under the sampled culture condition. The genome encodes complete glycolysis and pentose phosphate pathways, together with an almost complete tricarboxylic acid (TCA) cycle and expressed components of oxidative phosphorylation (fig S7, S8). This metabolic architecture is consistent with aerobic processing of soluble organic carbon after uptake. A complete glyoxylate shunt, provides metabolic routes for both carbohydrate and lipid-derived carbon. Fatty acid catabolism is supported by expressed β-oxidation enzymes predicted to localize to both peroxisomes and mitochondria based on subcellular targeting signals. The complete metabolic landscape is presented as pathway completeness scores in supplementary Fig. S7 and an interactive iPath3 pathway map in supplementary Fig. S8).

### A selectively expressed secretome encodes extracellular digestive capacity

To identify the extracellular machinery underlying osmotrophic nutrition, we predicted the secretome of *S. psychra* (Fig. 2, Table S3). The genome encodes 1,015 predicted secreted proteins, representing 5.75% of the proteome. Of these, 568 proteins received functional annotation, whereas 44% lacked recognizable conserved domains. The annotated enzymatic secretome comprised carbohydrate-active enzymes, proteases, lipases, and oxidoreductases (Table 1, S3). The remaining proteins include structural proteins, putative transport facilitators, and proteins of unknown function. Although predicted secreted proteins were not more highly expressed than non-secreted genes, specific enzymatic subsets of the secretome showed elevated expression (Fig. 3, Table S4, S5). Secreted carbohydrate-active enzymes (CAZymes) were expressed more strongly than non-secreted CAZymes and background genes, indicating targeted deployment of extracellular carbohydrate-degrading enzymes rather than broad upregulation of secretion. CAZymes were enriched approximately threefold among the top 10% most highly expressed genes (fig. S9). Secreted proteases dominate the most highly expressed secreted proteins, together with several abundant proteins lacking functional annotation (Table S3). This indicates that *S. psychra* does not simply maintain a broadly elevated secretion program, but instead selectively deploys a focused extracellular digestive machinery.

**Table 1:** Functional classes of secreted enzymes of *Saccharomycomorpha psychra*’s proteome and secretome and their protein similarity to fungal (FunSecKB2) and bacterial (SecretomeP 2.0) secretome databases; category ‘Both hit’ include fungal and bacterial hits.

| Functional class | No. in Proteome | No. in Secretome | Fungal hit | Bacterial hit | No Hit | Both Hit | % secretome | % proteome |
| --- | --- | --- | --- | --- | --- | --- | --- | --- |
| CAZymes | 109 | 64 | 44 | 11 | 20 | 11 | 6.30 | 0.62 |
| Proteases | 152 | 45 | 25 | 0 | 20 | 0 | 4.43 | 0.86 |
| Lipases | 96 | 21 | 4 | 0 | 17 | 0 | 2.07 | 0.54 |
| Oxidoreductase | 523 | 16 | 4 | 0 | 12 | 0 | 1.57 | 2.96 |
| Other Proteins | 16,800 | 870 | 80 | 2 | 780 | 8 | - | - |

**Figure 2:**
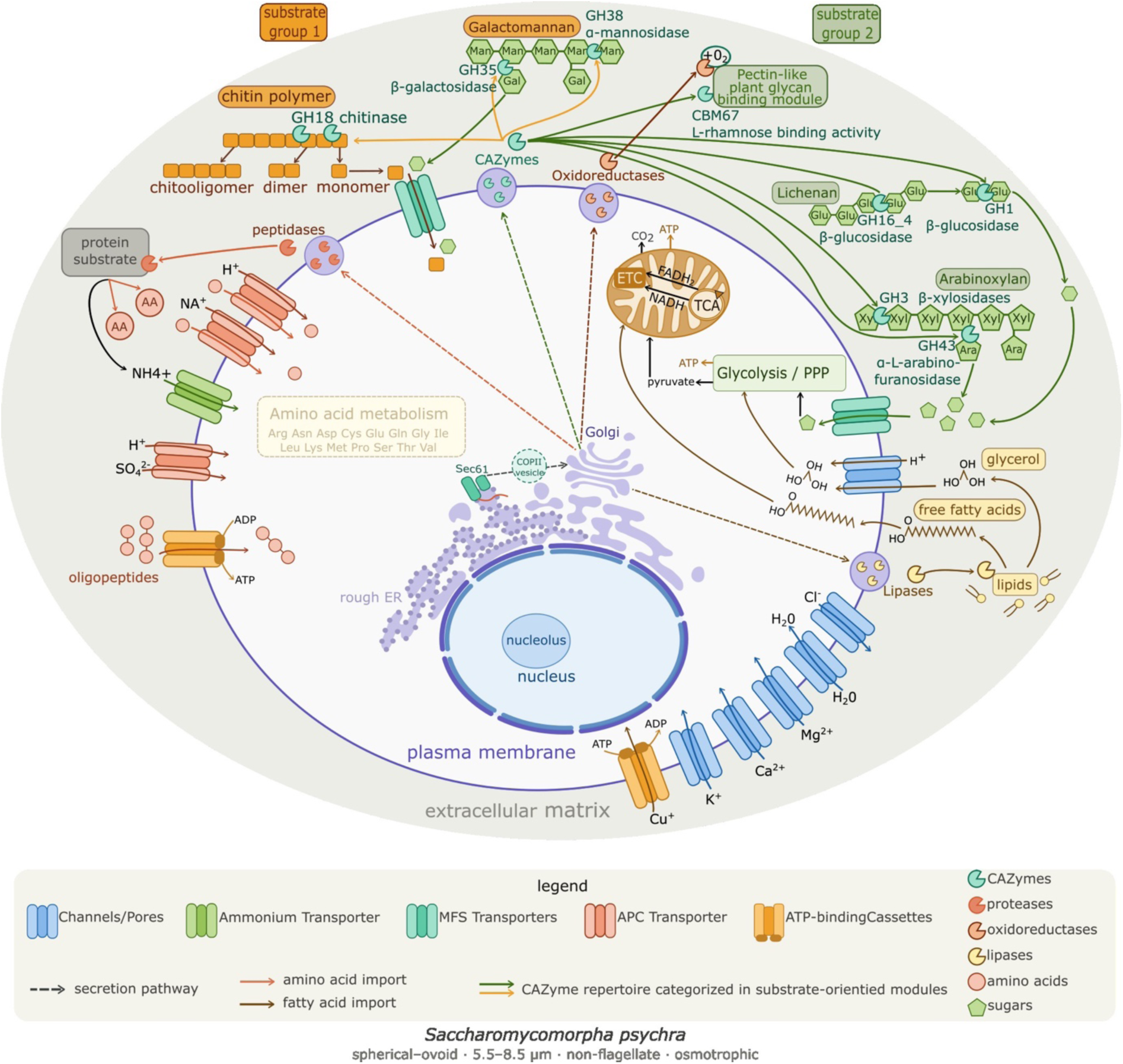
Genome-, transcriptome- and phenotype-informed reconstruction of osmotrophic metabolism in *Saccharomycomorpha psychra*. Schematic representation of the cellular organization and metabolic capabilities of *S. psychra* inferred from genome annotation, transcriptome profiling and carbon-utilization assays. Extracellular polymers associated with lichen-derived substrates are degraded by secreted carbohydrate-active enzymes (CAZymes), proteases, lipases and oxidoreductases. Resulting sugars, amino acids and lipids are imported through membrane transport systems, including major facilitator superfamily (MFS), amino acid–polyamine–organocation (APC) and ATP-binding cassette (ABC) transporters. Phenotypic assays confirmed utilization of multiple substrates, including D-/L-arabinose, L-rhamnose D-xylose and D-ribose, supporting the predicted capacity for uptake and metabolism of pentose-rich lichen-derived carbon sources. Imported carbohydrates are metabolized through glycolysis and the pentose phosphate pathway, with pyruvate feeding the tricarboxylic acid cycle and oxidative phosphorylation in mitochondria. Fatty acids enter β-oxidation in mitochondria, contributing acetyl-CoA to central carbon metabolism. Secreted proteins are synthesized and processed through the endoplasmic reticulum-Golgi secretory pathway before export to the extracellular space. Solid arrows indicate substrate degradation, metabolite transport and intracellular metabolic fluxes; dashed arrows indicate vesicle trafficking and protein secretion.

**Figure 3.**
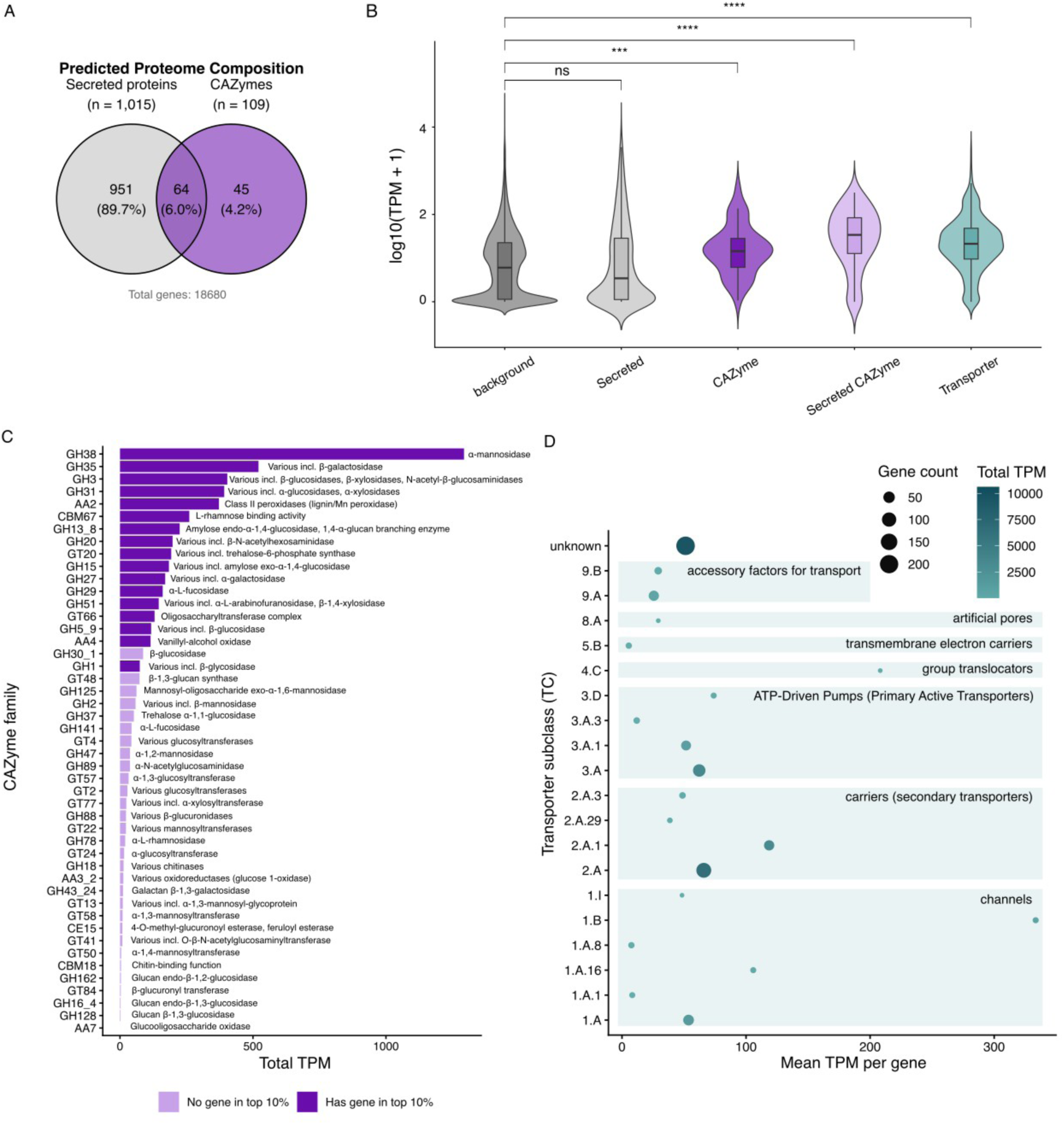
Secretome composition and transcriptional activity in *Saccharomycomorpha psychra*. **(A)** Overlap between predicted secreted proteins (n = 1,015) and carbohydrate-active enzymes (CAZymes; n = 109) within the predicted proteome (17,680 genes). The intersection comprises 64 secreted CAZymes. **(B)** Distribution of transcript abundance (log10[TPM + 1]) for all genes (background), secreted proteins, transporters, CAZymes and secreted CAZymes. Brackets indicate two-sided Wilcoxon rank-sum tests relative to the genomic background, with Benjamini–Hochberg correction for multiple testing. **(C)** Total transcript abundance summed across genes within each CAZyme family. Families are ordered by total TPM. Colors indicate whether at least one family member exceeds the transcriptome-wide 90th percentile expression threshold (69.4 TPM). **(D)** Abundance and expression of transporter subclasses. Bubble size indicates the number of genes assigned to each subclass, and color indicates total TPM summed across all genes within the subclass.

### Secreted CAZymes target fungal, lichen, and hemicellulose-derived substrates

CAZymes are classified into families designated as glycoside hydrolases (GH), auxiliary activities (AA), carbohydrate esterases (CE), glycosyltransferases (GT), and carbohydrate-binding modules (CBM), with family numbers indicating shared substrate specificity and catalytic mechanism. The CAZyme repertoire of *S. psychra* comprised 109 genes spanning 47 families, including 64 predicted secreted CAZymes (fig S10, Table S6, S7). This repertoire resolves into substrate-oriented modules (Table S8). The largest module targets fungal cell wall polymers and glycoproteins, accounting for 48% of total CAZyme transcript abundance (TPM). It encompasses enzymes that degrade chitin and β-glucans, the structural polysaccharides of fungal cell walls, alongside mannan-trimming and glycoprotein-remodeling activities (key families: GH18 chitinase, GH38/GH47/GH125 α-mannosidases, GH20/GH27 hexosaminidases and galactosidases, GH31 and GH5_9 α-/ and β-glucosidases; full family list in Table S7, S8).

A second module targets lichen polysaccharides and plant hemicellulose (27.9% of CAZyme TPM). This includes a GH16_4 lichenase, an enzyme specific to lichenan, the β-1,3/1,4-glucan backbone of the lichen thallus, alongside arabinose-releasing enzymes (GH43 arabinosidase, GH51 arabinofuranosidase), a β-galactosidase active on galacto-oligosaccharides (GH35), an α-rhamnosidase (GH78), and rhamnogalacturonan-targeting carbohydrate-binding modules (CBM67). A third, oxidative module included AA2-type class II peroxidase, AA3_2 glucose-methanol-choline oxidoreductase, AA4 vanillyl-alcohol oxidase, and CE15 glucuronoyl esterase, accounting for 8.9% of CAZyme TPM and indicating potential for oxidative modification of more recalcitrant aromatic or polysaccharide-associated substrates. With 3720 TPM, a single AA2-type class II peroxidase represents the most highly expressed individual CAZyme gene (Fig S10, Table S9). Consistent with the cofactor requirements of AA2 peroxidases, the genome encodes copper-importing metal ion channels (CopA-type). In contrast, canonical machinery for crystalline cellulose degradation was limited: GH6 and GH7 cellobiohydrolases were absent, and the annotated endoglucanase (K01179) was transcriptionally silent. The CAZyme profile therefore supports extracellular degradation enriched in activities targeting fungal glycans, glycoproteins of lichen-associated fungal and hemicellulose-rich substrates rather than a generalized plant-cellulose-degrading strategy.

### Proton-coupled transporters complete the osmotrophic uptake system

Extracellular digestion requires efficient import of the released soluble products. Consistent with this requirement, *S. psychra* encodes an expanded transporter repertoire comprising 680 putative membrane transporters (Table S10). Among TCDB-classified transporters, secondary active transporters (2.A; 112 genes) formed the largest class, followed by primary active transporters (3.A; 66 genes) and ion channels (1.A; 40 genes; Table S11). Major facilitator superfamily transporters represented the largest individual transporter group, consistent with proton-coupled uptake of dissolved organic compounds. APC Amino acid permeases and ABC transporters further support import of nitrogenous compounds and peptides. The presence of plasma membrane H⁺-ATPases provides a mechanism for generating the proton motive force required by the dominant MFS and APC transporter systems. Together, this transporter architecture, which is quantitatively dominated by proton-motive-force-dependent secondary carriers and significantly elevated in expression relative to background (Wilcoxon p = 2.7 × 10⁻^59^, Table S4), reflects the energetic demands of continuous active uptake of dissolved organic monomers from a dilute substrate, a hallmark of obligate osmotrophic physiology.

### Phenotypic assays validate broad pentose utilization

To test whether the predicted osmotrophic capacity translates into substrate use, we assayed carbon utilization using Biolog Phenotypic MicroArray**^TM^** PM1 and PM2, covering 192 carbon sources. *S. psychra* utilized 17 substrates, with the strongest responses observed for pentoses and modified hexose substrates (Fig S11, Table S12, S13). Notably, all six tested C5 monosaccharides were utilized, whereas C5 sugar alcohols produced no response, indicating that the capacity for C5 carbon uptake is specific to the free monosaccharide form rather than reflecting a general pentose affinity. Hereafter, “pentoses” refers specifically to pentose monosaccharides. This pentose utilization profile is consistent with the transcriptomic data: xylose isomerase (K01805) was highly expressed at 437.9 TPM, xylulokinase (K00854) at 36.5 TPM, ribose-5-phosphate isomerase (K01807) at 51.5 TPM, and ribokinase (K00852) at 35.5 TPM; all required for catabolism of pentoses. Together, these expressed enzymes provide direct metabolic entry points for xylose- and ribose-derived carbon into central metabolism. Utilization of D-arabinose despite the absence of a canonical arabinose isomerase suggests that arabinose-derived carbon may enter metabolism through an alternative route or through enzymes not resolved by current annotation. In contrast, several common hexoses and disaccharides, including D-glucose, showed little or no detectable utilization under the assay conditions. Thus, the phenotypic and transcriptomic data converge on a pentose-centered utilization profile, supporting the genomic prediction that *S. psychra* is adapted to dissolved carbon derived from pentose-rich lichen-associated substrates.

### The rhizarian secretome combines fungal-like enzymatic functions with extensive novelty

Comparative analysis of the predicted secretome revealed both similarity to fungal extracellular enzyme systems and a large uncharacterized, lineage-specific component. Across the full predicted secretome of 1,015 proteins, 18% showed detectable similarity to curated fungal secretomes, whereas only ∼2% matched bacterial secretome databases. This fungal similarity was concentrated in the enzyme classes most directly implicated in extracellular digestion: 44 of 64 secreted CAZymes showed fungal hits, and 25 of 45 secreted proteases showed fungal hits. In contrast, secreted lipases and oxidoreductases showed much weaker similarity, with only 4 of 21 lipases and 4 of 16 oxidoreductases matching fungal secretome entries. The remaining secretome was largely unresolved: approximately 80% of secreted proteins lacked detectable homologs in either fungal or bacterial secretome databases, including 20 secreted CAZymes, 20 secreted proteases, 17 lipases, 12 oxidoreductases, and 780 proteins outside these enzymatic classes. Thus, the *S. psychra* secretome combines recognizable fungal-like similarities among extracellular digestive enzymes, especially among CAZymes and proteases, with a substantial fraction of uncharacterized secreted proteins, underscoring both the functional analogy to fungal osmotrophy and the limited comparative resolution currently available for rhizarian extracellular biology.

## Discussion

### *Saccharomycomorpha psychra* as a model for free-living heterotrophic Rhizaria

*Saccharomycomorpha psychra* provides the first experimentally tractable genomic model for reconstructing heterotrophic metabolism in Rhizaria. Its osmotrophic specialization demonstrates how extracellular digestion can be organized within a supergroup whose metabolic biology remains poorly resolved despite its ecological abundance and diversity (Burki et al., 2021; Oliverio et al., 2020; Schoenle et al., 2025). By combining a telomere-anchored high-quality reference genome assembly, transcriptome-informed functional annotation, secretome and transporter prediction, and phenotypic carbon-utilization assays, we provide an integrated reconstruction of osmotrophy in a rhizarian protist.

The central finding is that *S. psychra* implements extracellular nutrition through a coupled secretion–uptake system: a selectively expressed secretome enriched in carbohydrate-active enzymes, proteases, lipases, and oxidoreductases depolymerizes complex substrates outside the cell, while an expanded transporter repertoire dominated by proton-coupled carriers imports the resulting soluble monomers. Carbon utilization profiling across 192 substrates using Biolog Phenotypic MicroArray**^TM^** confirmed that this predicted architecture translates into measurable substrate use, with 17 carbon sources actively utilized, including all six tested C5 monosaccharides. *S. psychra* is therefore not only a specialized lichen-associated protist, but also a tractable system for studying how extracellular digestion and dissolved-carbon uptake are organized within a major, poorly sampled eukaryotic supergroup.

### Osmotrophy as a derived specialization built on a rhizarian metabolic framework

In its broadest sense, osmotrophy denotes the uptake of dissolved substrates across the cell surface, but in organisms exploiting complex organic matter this requires prior extracellular depolymerization by secreted enzymes and subsequent uptake of the released low-molecular-weight products through membrane transporters (Richards & Talbot, 2018). This two-step principle is well established in fungal and stramenopile osmotrophs (labyrinthulomycetes and oomycetes), where secreted carbohydrate- and protein-degrading enzymes are coupled to transport systems that import the products of extracellular digestion (de Vries & de Vries, 2020; Liu et al., 2023; McGowan et al., 2020). *S. psychra* implements an analogous architecture in a rhizarian context. First, complex environmental polymers are depolymerized outside the cell by a selectively and differentially expressed secretome. Second, the released soluble products are imported through an expanded transporter repertoire in which MFS transporters, amino acid permeases, ABC transporters, and the plasma membrane H⁺-ATPase together provide the machinery for active uptake.

Nitrogen acquisition is integrated into the same process rather than running in parallel. *S. psychra* lacks dedicated pathways for inorganic nitrogen acquisition i.e. no nitrate reductase, nitrite reductase, nitrogenase, ammonia oxidation, dissimilatory nitrate reduction, or urea transporter was identified, indicating that inorganic nitrogen is unlikely to be a major nutritional input. Instead, nitrogen is derived primarily from extracellular proteolysis: a subtilisin-like serine protease and a cysteine-type peptidase are among the highest expressed genes, hydrolyzing environmental proteins into oligopeptides and amino acids for import. Ammonium released during this process, together with intracellular deamination products, is recovered by an Amt/MEP-family transporter (TC 1.A.11) and reassimilated via the strongly expressed GS/GOGAT pathway, which incorporates ammonium into glutamine and glutamate for biosynthesis. Carbon and nitrogen acquisition are therefore not separate systems but two arms of a single osmotrophic loop, extracellular digestion, proton-coupled uptake, ammonium recovery, and biosynthetic reassimilation, representing a reorganization of nutrient acquisition away from phagotrophic bacterivory and toward secretion-, uptake-, and respiration-centered metabolism.

### Lichen-associated substrate use and pentose-centered metabolism

Lichen thalli are structured microbial habitats in which fungal, photobiont, bacterial, and other eukaryotic partners contribute to a chemically heterogeneous matrix of cell-wall polymers, extracellular polysaccharides, secondary metabolites, and detrital organic matter (Grube et al., 2015). The CAZyme profile of *S. psychra* maps onto the principal biopolymer classes of the lichen microhabitat, rather than to general plant-cellulose degradation.

The dominant module targets the chitin, β-1,3-glucan, and galactomannan-rich glycoprotein matrix of the lichen mycobiont cell wall (48% of CAZyme TPM), and a second module targets lichenan and hemicellulosic polysaccharides of the lichen thallus and associated plant debris (27.9% of CAZyme TPM). Together these two ecologically interpretable modules account for 76% of CAZyme transcript output. The absence of cellobiohydrolases and transcriptional silencing of the single annotated endoglucanase argue against crystalline cellulose as a major substrate. The phenotypic carbon-utilization profile validates this interpretation. All six tested pentoses were utilized, matching the high expression of pentose catabolic enzymes and consistent with the predicted degradation of arabinose-, xylose-, galactose-, and rhamnose-containing polysaccharides by the secreted CAZyme repertoire. Conversely, D-glucose showed little or no detectable utilization, despite encoding complete glycolysis, suggesting either a regulatory or transport-level limitation whose basis remains unresolved. Lichen mycobionts cycle hexoses for their own energy metabolism and export photobiont-fixed carbon predominantly as the sugar alcohols ribitol and arabitol (Fahselt, 1994); a protist specialized on the pentoses released from hemicellulosic side chains within the same thallus would therefore occupy a chemically distinct carbon niche, consistent with resource partitioning rather than direct competition. The convergence of mannan-targeting CAZyme expansions across lichen-associated bacteria, lichen-forming fungi, and a rhizarian protist, three phylogenetically independent lineages sharing the same microhabitat, suggests that galactomannan is a recurrent target of carbon acquisition within lichen thalli.

### A convergent fungal-like osmotrophic architecture

The functional parallel of *S. psychra* secretome and those of saprotrophic fungi extends beyond individual gene families to overall architectural organization: 44 of 64 secreted CAZymes and 25 of 45 secreted proteases show detectable similarity to fungal secretome entries (68% of secreted CAZymes carrying fungal homologs), whereas bacterial secretomes similarity was limited. Lichen-forming fungi (Lecanoromycetes) retain extensive CAZyme arsenals despite stable photosymbiosis, including expansion of GH5 1,3-β-glucosidases (Resl et al., 2022), the same subfamily present in *S. psychra*, indicating that both organisms are independently equipped to act on mycobiont β-1,3-glucan within the same microhabitat. Likewise, lichen-associated bacterial communities have additionally documented an independent expansion of GH38, GH92, and GH125 mannan-targeting enzymes in Acidobacteriaceae (Tagirdzhanova et al., 2024), indicating that galactomannan constitutes a carbon resource targeted by multiple phylogenetically unrelated lineages sharing this microhabitat. The expansion of mannan processing CAZyme families in bacteria, lichen-forming Lecanoromycetes, and now a rhizarian protist, points to the galactomannan matrix of the mycobiont cell wall as a convergently exploited resource within lichen ecosystems. Beyond extracellular enzymes, *S. psychra* shares the canonical osmotrophic architecture of fungi, combining a large secretome with H⁺-coupled MFS transporters, which dominate sugar import alongside proton-motive-force generation by the plasma membrane H⁺-ATPase (Rottmann et al., 2018)*. S. psychra* additionally encodes and shows transcriptional activity of an AA2-type class II peroxidase that is restricted to Basidiomycetes and some Ascomycetes within fungi. However, as it lacks a detectable secretion signal, its subcellular localization and functional role within the cell remain to be characterized.

Whether the architectural similarity reflects convergent recruitment, ancient retention with lineage-specific expansion, or horizontal gene transfer from lichen-associated Ascomycota or Basidiomycota cannot be resolved from annotation alone. The lichen microhabitat provides sustained physical proximity between *S. psychra* and fungal hyphae, a setting documented to facilitate inter-kingdom gene transfer in microbial eukaryotes (Keeling & Palmer, 2008; López-García & Moreira, 2026). Conversely, many of the relevant CAZyme families are broadly distributed across eukaryotes, making repeated functional recruitment equally plausible. Resolving these alternatives will require phylogenetic reconstruction of individual gene families.

The only phagotrophic rhizarian with transcriptomic characterization of CAZyme deployment, *Orciraptor agilis*, expresses a toolkit specialized for algal cell-wall perforation dominated by GH5_5 cellulases, GH18 chitinases and pectinases (Gerbracht et al., 2022), lacking the mannan-targeting, lichen-associated and oxidative modules characteristic of *S. psychra*. Conversely, osmotrophic oomycetes converge on the same secretion–uptake strategy but deploy cellulases and polygalacturonases adapted to plant cell walls. *S. psychra* therefore represents a distinct, lichen-adapted implementation of a broadly convergent osmotrophic architecture.

Regardless of its evolutionary origin, the architectural similarity between *S. psychra* and fungal osmotrophs spans one of the deepest evolutionary divisions among extant eukaryotes. Molecular clock analyses place the divergence of SAR, including Rhizaria, and Opisthokonta, including fungi, deep in the Proterozoic, more than one billion years before present (Strassert et al., 2021). Crown-group fungi diversified substantially later (Szánthó et al., 2025). The emergence of a functionally similar osmotrophic system across this evolutionary distance suggests that comparable ecological pressures can repeatedly favor the assembly of analogous extracellular digestive and nutrient uptake architectures.

### Lineage-specific proteins and the limits of comparative genomics

Approximately 44% of predicted secreted proteins lack recognizable conserved domains, and around 80% lack detectable similarity to the fungal or bacterial secretome databases queried here. This unresolved fraction should not be read as annotation failure: secretomes of fungi and other microbial eukaryotes characteristically contain disproportionate numbers of lineage-specific or poorly characterized proteins relative to the cytoplasmic proteome, including small secreted proteins whose functions cannot be inferred from sequence similarity alone (Haas et al., 2009; Pellegrin et al., 2015; Stergiopoulos & de Wit, 2009). This limitation is amplified in Rhizaria, where genome resources remain sparse and taxonomically biased (Schoenle et al., 2025). The large unannotated fraction thus most plausibly reflects a combination of genuine rhizarian novelty and the current limits of comparative annotation, a distinction resolvable only through expanded rhizarian genome sampling, structural prediction, and substrate-specific transcriptomics. The subset of secreted proteins showing low or absent expression under the single axenic culture condition sampled here likely constitutes an inducible substrate-responsive layer, analogous to the inducible CAZyme component documented in saprotrophic fungi (Znameroski & Glass, 2013), rather than non-functional sequence. *S. psychra* as a genomically tractable model system amenable to culture, transcriptomic profiling across defined substrates, and phenotypic assay, is ideally placed to resolve both the lineage-specific and inducible components of this hidden secretome.

### Trophic transitions in Rhizaria and the wider landscape of eukaryotic osmotrophy

The reconstruction of osmotrophy in *S. psychra* also provides a framework for studying trophic transitions within Rhizaria. Most characterized direct relatives are phagotrophic bacterivores, whereas several well-studied rhizarians are parasitic or endobiotic, for example, the clubroot pathogen *Plasmodiophora brassicae*, whose genome analyses reflects metabolic features shaped by obligate biotrophy (P. Li et al., 2025; Pérez-López et al., 2020; Schwelm et al., 2015). *S. psychra* occupies a third functional position: free-living, cultivable, non-phagotrophic, and equipped for extracellular digestion of environmental polymers, with a secretome size of 5.75% of the proteome. This three-way contrast, phagotrophic bacterivore, biotrophic parasite, and free-living osmotroph, suggests that osmotrophy in Rhizaria arose not through wholesale replacement of central metabolism but through reorganization of the cell surface and extracellular interface around an otherwise conventional aerobic heterotrophic core: selective expansion and deployment of secreted enzymes, transporters, and proton-motive uptake systems.

Within the wider landscape of eukaryotic osmotrophy, *S. psychra* represents an unusually complete case of polysaccharide-based extracellular digestion in a protist. Coccolithophores, certain phytoflagellates, and heterotrophic nanoflagellates, scavenge simple sugars and amino acids via osmotrophic uptake (Balch et al., 2023). Among Amoebozoa, *Dictyostelium* supplements phagotrophy with macropinocytic uptake of dissolved protein (Lutton et al., 2023; Rivero & Maniak, 2006). Comparable polymer-degrading osmotrophic strategies are found in fungi-like Labyrinthulomycetes and oomycetes, whose genomes encode expanded GH repertoires and secreted lipases (Liu et al., 2023; McGowan & Fitzpatrick, 2020). However, the enzymatic profile of *S. psychra* differs markedly from that of a plant-pathogenic oomycete *Phytophthora,* which encodes expanded hemicellulases (GH10) and pectinases (GH28, PL) and crystalline-cellulose-targeting cellulases (GH6/GH7) adapted for a host-cell wall penetration during infection (Zerillo et al., 2013). The absence of these plant-perforation toolkit in *S. psychra,* together with its enrichment in fungal cell-wall and glycoprotein-targeting enzymes, reflects adaptation to a free-living decomposition of lichen-associated biopolymers. Comparative genomic sampling across phagotrophic, parasitic, and osmotrophic Rhizaria, currently limited by the scarcity of reference genomes, will be required to determine which components of this osmotrophic system are ancestral, which are lineage-specific expansions, and which are mechanistically tied to the loss of phagotrophy.

## Conclusion

We present the first integrated genomic, transcriptomic, and phenotypic reconstruction of an osmotrophic lifestyle within Rhizaria, grounded in a high-quality telomere-anchored reference genome for a free-living member of this underrepresented supergroup. *S. psychra* has assembled a secretion-based extracellular digestive system, a hemicellulase and glycoprotein-targeting CAZyme repertoire mapped onto lichen-associated substrates, coupled to a proton-driven transporter arsenal, in an organism whose immediate relatives rely exclusively on phagotrophic bacterivory. The experimental confirmation of broad pentose utilization validates the CAZyme- and transporter-based predictions and closes the loop between genome, transcriptome, and phenotype. These findings indicate that shared ecological pressures, specifically, the demand for complex polysaccharide decomposition within a lichen-dominated polar microhabitat, can drive convergent enzymatic and transport solutions across vast phylogenetic distances, independent of shared ancestry. *S. psychra* thereby provides both a tractable experimental platform for studying the genomic and mechanistic basis of osmotrophic lifestyle transitions in protists, and an entry point for testing how trophic transitions, secretome evolution, and extracellular nutrient acquisition have shaped the ecological diversification of Rhizaria.

## Methods

### Sample preparation and sequencing

#### Sample collection

Samples were collected from various types of moss, and lichen at Byers Peninsula, Antarctica [62°39’54.7”S 61°06’13.6”W] in January 2023. Samples were stored and transported at -80°C before use. Following Feng et al. 2021, samples i.e. biocrusts and lichen were cut into pieces of one to two centimeters and immersed in 75% EtOH (Merck, Germany) for one minute, followed by 1% sodium hypochlorite (VWR, Germany) for 30 seconds and 75% EtOH for 30 seconds. Then pieces were washed in distilled water and dried on sterile filter paper (Satorius Biolab) and manually screened for *Saccharomycomorpha psychra*.

#### Sample preparation

For cultivation 39 g L-1 of powdered Millipore Potato Glucose Agar (PGA) was prepared with distilled water. After sterilization in the autoclave, doxycycline (Cayman Chemical, Hamburg, Germany) and streptomycin (Sigma Life Science, Darmstadt, Germany) were added to the cooled-off liquid agar, each up to a concentration of 50 mg L-1. The agar was poured into Petri dishes (VWR). The prepared pieces are evenly spread and sealed with Parafilm (Bemis). The plates with samples were stored at 15 °C.

#### Species identification

Putative *Saccharomycomorpha* colonies were identified based on morphology and established as monoclonal cultures. DNA was extracted using a modified guanidine-based protocol following Pawlowski et al. (2000). The 18S rRNA gene was amplified using universal eukaryotic primers EukA and 1300r. Amplicons were confirmed by agarose gel electrophoresis and bidirectionally Sanger sequenced (Eurofins Genomics). Consensus sequences were assembled from forward and reverse reads, compared to NCBI reference sequences using BLAST, aligned with representative Cercozoa sequences using MAFFT, and analyzed by maximum-likelihood phylogenetic inference in RAxML (GTR+I+G; 200 bootstrap replicates). Strain 255 was recovered as the closest relative of the *S. psychra* type strain and was therefore selected for genome sequencing and all downstream analyses.

### High-molecular-weight *g*DNA extraction

To isolate high-molecular-weight (HMW) *g*DNA of *S. psychra* strain 255, a modified version of Öztoprak & Bast, 2023 was used. In short: clonal specimen from an agar plate were scraped off and distributed among 3 Eppendorf tubes. Each tube was submerged in 380 µl TNES buffer and flash-frozen three times in liquid nitrogen. Proteinase K was added to the homogenized sample and incubated for 1h at 37°C. RNase cocktail was added and incubated at 37°C for 30 min. Then, NaCl and absolute ethanol were added, and the sample was incubated for at least 1h at -20°C. DNA was purified and sample homogenized at 4°C overnight. DNA concentration was measured using Qubit Fluorometer v.4 with the Qubit dsDNA HS Assay Kit (Thermo Fisher Scientific, Waltham, MA).

DNA quality was assessed at the Cologne Center of Genomics (CCG, Cologne, Germany) using Genomic DNA ScreenTape®. HMW *g*DNA with ∼3 µg of DNA, fragment size above 15 kb and DIN > 8.0 was selected for sequencing.

### Long-read sequencing for *de novo* assembly

The PacBio HiFi SMRTbell® Library for whole genome sequencing was used. Library preparation and sequencing of the SMRTbell^TM^ templates were conducted by the Genomics & Transcriptomics Labor (Düsseldorf, Germany). Sequencing yielded in total 14.2 Gb of Pacific Biosciences Circular Consensus HiFi reads (Wenger et al. 2019) with an N_50_ of 9.4 kb.

#### RNA sequencing

To isolate total RNA, specimen were once washed with Sorenson buffer, then RIN-buffer was used to flash-freeze the sample (three times) with liquid nitrogen. Next, the RNeasy Plant Mini Kit for RNA Extraction (Qiagen) was followed. RNA quality was assessed on a TapeStation (CCG). To construct the RNA-seq library the CCG used the Illumina TruSeq Stranded mRNA kit for bulk mRNA sequencing of 100-bp reads.

#### Assessing HiFi reads

To judge sequencing complexity such as ploidy, heterozygosity and genome duplications we conducted *k*-mer analyses on HiFi reads of *S. psychra*. 27-mers in the HiFi reads were analyzed using KAT v2.4.2 (Mapleson et al., 2017) with the modules kat hist and kat gcp (default parameters). Ploidy was further investigated using kmc v3.2.1 with parameters -k 27 -ci 1 -cs 10000 and Smudgeplot v0.2.5 (Ranallo-Benavidez et al., 2020) with default parameters. To obtain a frequency count of *k*-mers Jellyfish v2.2.8 (Marçais & Kingsford, 2011) was used as input to estimate genome heterozygosity and repeat content with GenomeScope2.0 (http://genomescope.org/genomescope2.0/; last access 7.07.26; Ranallo-Benavidez et al., 2020).

### Assembly and assembly evaluation

#### *De novo* assembly

HiFi reads were assembled using hifiasm v0.25.0 (Cheng et al., 2021) with parameter ‘-l 0’. Contigs were manually curated using BlobTools2 v4.3.2 (Challis et al., 2020), including identifying contaminants and assembling homozygous regions of high coverage, which were not fully resolved, through the graphical assembly.

HiFi reads were additionally assembled with PECAT (Nie et al., 2024) as it includes a haplotype-aware correction method. The PECAT assembly was scaffolded with the curated hifiasm assembly as reference using RagTag v2.1.0. (Alonge et al., 2022).

#### Completeness

To assess ortholog completeness Benchmarking Universal Single-Copy Orthologs (*BUSCO* v5.0.0; Manni et al., 2021; Simão et al., 2015) was used against the Eukaryota odb10 lineage (255 orthologs), Alveolata odb 10 lineage (171 orthologs). To evaluate *k*-mer completeness, KAT v2.4.2 was used with the module kat comp with default parameters. To identify potential contamination mapped HiFi reads were mapped to the final scaffolds using minimap2 v2.24 (H. Li, 2021) with parameters ‘-ax map-hifi’ and the mapped reads were sorted with SAMtools v1.11. The final scaffolds were aligned against the nucleotide database using the Basic Local Alignment Search Tool (BLAST) v2.6.0 (Altschul et al., 1990) with parameters ‘-outfmt “6 qseqid staxids bitscore std sscinames scomnames” -max_hsps 1 - evalue 1e-25’. The outputs of minimap2, BLAST, and BUSCO (against the Eukaryota odb10 lineage) were provided as input to BlobTools2. Sequences identified as bacteria were subsequently removed.

### Genome annotation

#### Repeat and transposable element annotation and masking

Transposable elements (TE) and repetitive regions were predicted with the Extensive *De novo* Annotator (EDTA) pipeline v2.2.2 (Bell et al., 2022) with parameters ‘—sensitive 1 –anno 1’. RepeatModeler v2.0.1 (Flynn et al. 2020) was used to identify repeat element boundaries of the TE library. TEs were classified using the FasTE pipeline (Bell et al., 2022) to superfamily level using the convolutional neural network program DeepTE (python = v3.6; tensorflow-gpu = v1.14.0; biopython; keras = v2.2.4; numpy = v1.16.0; Yan et al., 2020). To annotate TEs against the custom library, we used RepeatMasker v4.2.2 (Tarailo-Graovac & Chen, 2009) with parameters ‘-a -s -no_is’. Prior to downstream analysis, the output files were processed and filtered with the script “RM_Trips” in R (v4.4.3; R Core Team, 2025). Telomeric repeat candidates were identified using the telomere identification toolkit tidk v0.2.41 (Brown et al., 2025). Each scaffold was subsequently classified according to the presence and position of telomeric repeat arrays at its termini using a custom Python pipeline: scaffolds with telomeric repeats detected within 20 kb of both ends were classified as ’complete’ (chromosome-level), scaffolds with a telomeric repeat at only one terminus as ’single-telomere’, and scaffolds lacking detectable telomeric repeats at either terminus as ’no-telomere’. This classification was used to define the 22 chromosome-level scaffolds (complete) and the additional 26 single-telomere scaffolds reported.

The hardmasked assembly was converted into a softmasked assembly using bedtools (v2.30.0). Superfamilies were renamed according to Wicker-classification (Wicker et al., 2007). Divergence landscapes of TE superfamilies were generated via custom R Scripts described in Öztoprak et al., 2025.

### *De novo* annotation

RNA-seq reads were trimmed using FastP v0.21.0 with parameters ‘-q 20 -u 5 -l 25 -y -n 0’ and mapped to the assembly using hisat2 v2.2.1 (Kim et al., 2019). Mapped reads were sorted using Samtools v1.6 (Danecek et al., 2021). For annotation BRAKER v3.0.8 (Gabriel et al., 2024), with parameters ‘--gff3 --UTR off’ was used, which combined predictions from Augustus v3.5.0 and GeneMark-ES v4.68. The longest isoform was selected using Another Gtf/Gff Analysis Toolkit (AGAT) v0.8.0. Completeness of predicted proteins was evaluated using BUSCO v5.4.7 against the Eukaryota odb10 and Alveolata odb10 lineages with the parameter ‘-m protein’.

#### Genome landscape visualization

Sequencing coverage, GC content, and gene, exon, and transposable-element density were calculated in non-overlapping 100-kb windows. All tracks were visualized as geom_rect layers to ensure consistent x-axis alignment across panels, using ggplot2 v4.0.2 and patchwork v1.3.2 in R v4.5.1. Individual scaffold panels were combined using wrap_elements(full = row) v1.3.2 to preserve relative panel proportions. Figures were rendered using the svglite v2.2.2. and ragg v1.5.0 graphics devices.

#### Functional annotation

To functionally categorize the protein sequences Eggnog-mapper v2.1.4 (Cantalapiedra et al., 2021) with parameters ‘--score 0.01 --seed_ortholog_evalue 0.01’ and InterProScan v5.69-101.0 (Jones et al., 2014) were employed.

Carbohydrate-activated-enzymes (CAZymes) were predicted by multiple evidence layers from Hmmer, Hotpep and DIAMOND by dbCAN2 (Zhang et al., 2018). Only proteins supported by at least two independent annotation approaches were retained, yielding 109 CAZymes across 47 families.

To further characterize essential metabolic redox reactions enzymatic functional groups i.e oxidoreductases based on InterPro domains associated with oxidoreductase activity; proteases using EC 3.4.x.x assignments, MEROPS HMM searches ‘with an e-value threshold of 1e−5 hits were filtered to retain alignments with ≥30% amino acid identity and ≥50% query coverage’, and InterPro peptidase domains; lipases as proteins with EC 3.1.1.x and domain-based belonging to known lipase PFAM families (PF00151, PF01764, PF00657), were identified. All functional categories were quantified both relative to the total proteome and relative to the predicted secretome.

Putative transporter proteins were identified by combining transmembrane helix prediction via DeepTMHMM (Hallgren et al., 2022), a deep learning-based topology predictor, homology searches against the Transporter Classification Database (TCDB; Saier et al., 2021), EggNOG transporter classification, and PFAM domain annotation. Proteins containing at least one predicted transmembrane helix and at least one transporter-related annotation were retained as candidates. All evidence sources were integrated and proteins were assigned to one of three confidence tiers: high-confidence transporters were defined as proteins containing ≥3 evidence lines, medium confidence (n = 2), indicating agreement between two independent sources and low confidence (n =1 with ≥1 predicted TM helix), indicating a single transport-specific annotation supported by membrane topology.

Identified transporters were functionally classified according to the TC system hierarchy following class 1 (channels/pores), class 2 (secondary active carries), class 3 (primary active pump), class 4 (group translocators), class 5 (transmembrane electron carriers) and class 9 (incompletely characterized transporters). For genes lacking TC assignment (’unknown’), TC class was resolved by Pfam domain mapping using a curated lookup Table derived from TCDB domain annotations. The remaining 28.6% lacked TC assignment due to absence of diagnostic Pfam domains or highly divergent sequences. In total, 680 transporter-encoding genes were identified, of which 303 were of high-confidence.

#### Functional motifs in secreted enzymes

To identify secreted enzymes, we employed a comprehensive *in silico* pipeline integrating multiple prediction tools. SignalP 6.0 was used to detect classical signal peptides (SPs) and their cleavage sites, providing high-confidence predictions across all domains of life. To mitigate the common issue of cross-prediction between signal peptides and transmembrane regions, we additionally used Phobius (https://phobius.sbc.su.se, last accessed 23.03.2026), which combines SP detection and transmembrane topology prediction using a hidden Markov model (HMM). Since the presence of signal peptide alone does not determine protein localization, we incorporated DeepLoc 2.1 (Ødum et al., 2024) a multi-label predictor, based on machine-learning predicting eukaryotic protein subcellular localization and membrane associations. To further refine our secretome predictions, we assessed glycosylphosphatidylinositol (GPI) anchoring using NetGPI (Gíslason et al., 2021), which identifies post-translational membrane tethering via GPI anchors, which are frequently associated with non-secreted membrane-bound proteins.

To cross-validate all secreted enzyme candidates, we developed an inhouse filtering protocol that integrates the outputs of the above tools. This workflow resolves signal peptide and transmembrane conflict, filters GPI-anchored proteins, and flags competing subcellular localization assignments (e.g., mitochondrial, nuclear, or cytoplasmic predictions). This yielded 1,015 candidate secreted proteins.

#### Secretome comparison

To identify homologs of predicted secreted proteins across major microbial lineages, we performed comparative analyses against curated secretomes from fungi, and bacteria. These comparisons were conducted using FunSecKB2 including 167 species of fungi (Meinken et al., 2014) and SecretomeP 2.0 (Bendtsen et al., 2004). Hits with ‘DIAMOND (E ≤ 1e−5, ≥30% identity, ≥50% coverage)’ were interpreted as secreted proteins as described in other eukaryotic systems. Absence of detectable homology was considered indicative of potentially lineage-specific proteins.

#### Expression analysis

All analyses were performed in R (v4.5.1) using the tidyverse, ggplot2 v4.0.2, dplyr v1.2.0, rstatix v0.7.3, ggpubr v0.6.2, patchwork v1.3.2, and ggvenn v0.1.19 packages. RNA-seq expression data were obtained from Salmon (Patro et al., 2017) quantification output. For genes with multiple annotated isoforms, expression was summarized at gene level by retaining the isoform with the highest TPM value, yielding one expression estimate per gene (17,066 genes total). Expression analyses were performed only on genes recovered in the Salmon quantification matrix, whereas genome-wide annotation statistics refer to the complete predicted gene set. To stabilize variance and avoid undefined values for zero-expressed genes TPM values were log₁₀-transformed after addition of a pseudocount of 1 [log₁₀(TPM + 1)].

#### Statistical Comparisons of Expression

Genes were classified into functional categories (i) secreted CAZyme [secreted AND CAZyme], (ii) CAZyme [CAZyme only], (iii) transporter, (iv) secreted [secreted only], (v) background [all remaining genes]. Expression distributions were compared using Wilcoxon rank-sum tests. This non-parametric test was selected because TPM values were non-normally distributed with extreme right skew, the data were zero-inflated (∼18% of background genes had TPM = 0), and group variances were unequal. For each functional category, expression levels were compared against all genes not belonging to that category (background). All pairwise comparisons were corrected for multiple testing using the Benjamini-Hochberg (BH) false discovery rate procedure. Effect sizes were reported as rank-biserial correlations (r) using wilcox_effsize(), interpreted following Cohen (1988): |r| ≈ 0.1 small, 0.3 medium, 0.5 large.

#### Enrichment Analysis

The most highly expressed families were identified based on total TPM, assessed using Fisher’s exact test. To assess whether CAZymes are overrepresented among highly expressed genes, genes in the top 10% by TPM was defined using the gene-level 90th percentile threshold (TPM ≥ 69.4). A family was classified as ‘highly expressed’ if at least one of its member genes exceeded the threshold. Expression distributions were visualized using violin and boxplots. CAZyme family expression patterns were visualized using bar plots and heatmaps based on log-transformed TPM values.

#### Length-Bias Assessment

As a quality control measure, Spearman’s rank correlation was computed between gene length to verify that TPM normalization eliminated any residual length bias.

### Metabolic pathway reconstruction

To assess core biosynthetic pathways of microbial ecology and biochemical cycling, excluding less-conserved pathways KEGGaNOG v1.1.19 (Popov et al., 2026) was used, translating EggNOG annotations into KEGG module completeness scores. For full metabolic pathway reconstruction KofamKOALA (https://www.genome.jp/tools/kofamkoala/; ver. 2026-03-01; last accessed on 31.03.2026) was used. KofamKOALA allows for the annotation of diverged proteins as it enables KEGG Ortholog (KO) annotations, based on HMMER/HMMSEARCH searches against the customized KOfam database of KEGG Ortholog profiles (Aramaki et al., 2020). The annotation with standard settings (e-value 0.01), resulted in 19,7 % annotations. To visualize present pathways and their expression in a map iPath3 (; last accessed 31.03.2026) was used. Interactive pathways can be accessed via iPath3 with input data provided at https://github.com/HuesnaOeztoprak/Saccharomycomorpha.

### Experimental Carbon utilization profiles

#### Biolog Phenotypic MicroArray^TM^ Assays (PM1 and PM2)

Biolog Phenotypic MicroArray^TM^ plates PM1 and PM2 were used to assess the utilization of different carbon sources by *S. psychra* under controlled *in vitro* conditions.

*S. psychra* was cultivated on potato dextrose agar (PDA; 15 g/L agar, 20 g/L dextrose, 4 g/L potato extract) at 15 °C for three weeks. Biomass from approximately one PDA plate was aseptically collected using a sterile spatula and transferred into IFY-0 medium (1.2×; Biolog Inc., Hayward, CA, USA) supplemented with 500 µL/L Streptomycin (Sigma Life Science) and 75 µL/L Doxycycline (Cayman Chemical Company, Ann Arbor, MI, USA).

To obtain a homogeneous cell suspension, aggregated cells were gently disrupted by manual pestling and repeated pipetting. The optical density of the cell suspension was adjusted to OD₆₀₀ = 0.6 using a photometer (Thermo Scientific Varioskan, Waltham, MA).

PM additive solutions were prepared according to the manufacturer’s Phenotypic MicroArray**^TM^** Protocol for Yeast (Biolog Inc.), which was provided upon request. For PM plates 1 and 2, a total of 24 mL inoculation fluid was prepared, consisting of 20 mL IFY-0 (1.2×) medium (Biolog Inc.), 240 µL Redox Dye Mix D (Biolog Inc.), 500 µL cell suspension, 3.246 mL sterile water, 12 µL streptomycin and 1.8 µL doxycycline. Subsequently, 100 µL of the inoculation mixture was dispensed into each well of the PM plates.

The plates were incubated at 8 °C for a total period of 67 days with the lid closed. Metabolic activity was monitored by time-course imaging of the colorimetric readout: reduction of the tetrazolium redox dye to purple formazan by respiratory activity in the presence of utilizable substrate, following the manufacturer’s assay design. A substrate was classified as utilized if it produced a visually unambiguous and sustained color change (colorless to purple/violet) at any point during incubation. In addition, absorbance measurements were recorded at wavelengths 590 nm on day 0 and day 67.

## Acknowledgments

We thank Dr. Jule Freudenthal for valuable scientific input and discussions that contributed to the conceptual development of this work. We additionally thank the Cologne Center for Genomics (CCG, Cologne, Germany) and Genomics & Transcriptomics Laboratory (GTL; Düsseldorf, Germany) for their sequencing service and continuous support.

## Author contributions

K.D. and F.K. collected samples, H.Ö. performed genome assembly, annotation, and bioinformatic analyses. J.G. contributed to metabolic pathway reconstruction and transposable element analyses. R.J. contributed to the phenotypic profiling. K.D. supervised the project and contributed to data interpretation. H.Ö. wrote the manuscript with input from all authors. All authors read and approved the final manuscript.

## Data and materials availability

Sample collection in Antarctica was carried out in compliance with the Protocol on Environmental Protection to the Antarctic Treaty under permit II 2.3–94033/216 (Umweltbundesamt, Germany, issued 23 November 2022). Cultures of *Saccharomycomorpha psychra* strain 255 are deposited at the German Collection of Microorganisms and Cell Cultures (DSMZ, Braunschweig, Germany) under accession number [*to be inserted once submission is completed; current status: submitted*] and are available upon request. All code and scripts used for genome assembly, annotation, and downstream analyses are available at https://github.com/HuesnaOeztoprak/Saccharomycomorpha and have been archived with a citable DOI on Zenodo (10.5281/zenodo.21397006). Raw PacBio HiFi and RNA-seq reads as well as genome assembly are deposited at NCBI SRA under BioProject: PRJNA1491632.

