## Supplementary Material, Note and Figures for "A rhizarian genome reveals an osmotrophic route to extracellular digestion in eukaryotes"

Hüsna Öztoprak et al.

*corresponding author:

Hüsna Öztoprak,

Kenneth Dumack,

This includes:

Supplementary Text

Figs S1 to S11

Tables S1 to S13

#Data S1 to S3

**Supplementary text**

Supplementary Note to results section 2: Central metabolism supports aerobic processing of imported dissolved carbon

KEGG annotation identified a complete glycolytic pathway, with all enzymes transcriptionally detected and expressed under the sampled culture condition (Fig. S7, S8). The pentose phosphate pathway was complete. Components of oxidative phosphorylation were also detected, including subunits of NADH:ubiquinone oxidoreductase, ubiquinol-cytochrome c reductase (K00411), and V-type ATPase complexes (K02145–K02151). Canonical nitrate reductase (K00370) and 2-oxoglutarate ferredoxin oxidoreductase (K00174) were not detected. Several pathways involved in sugar utilization were recovered. The Leloir pathway was complete, including galactokinase (K00849), galactose-1-phosphate uridyltransferase (K01785), and UDP-galactose 4-epimerase (K01784), all of which were transcriptionally detected.

Genes encoding all amino acid biosynthetic pathways annotated in KEGG were detected. Components of the glutamine synthetase/glutamate synthase (GS/GOGAT) system were present and expressed, including glutamine synthetase (K01915; three gene copies) and glutamate synthase (K00264). Aminotransferases and aminopeptidases associated with amino acid turnover were also detected. Thus, rather than relying on independent inorganic nitrogen assimilation pathways, *S. psychra* appears to operate a tightly coupled osmotrophic nitrogen economy in which extracellular digestion, nutrient uptake, ammonium recovery, and GS/GOGAT-mediated reassimilation form an integrated recycling system.

Genes associated with fatty acid metabolism included 3-oxoacyl-ACP reductase (K00059) and acyl carrier protein (K00645). Enzymes involved in β-oxidation, including enoyl-CoA hydratase (K01692) and 3-hydroxyacyl-CoA dehydrogenase (K00022), were detected and transcriptionally expressed.

Together, these data support a metabolically versatile aerobic lifestyle in which imported carbohydrates are channeled into central carbon metabolism, oxidative phosphorylation and anabolic biosynthesis.

Supplementary Note to results section 5: Proton-coupled transporters complete the osmotrophic uptake system

The transporter repertoire of *Saccharomycomorpha psychra* was predicted using a multi-evidence framework integrating TCDB annotation, Pfam domain assignments, and transmembrane helix predictions (Table S10). This approach identified 724 putative membrane transport-associated proteins, corresponding to 3.85% of the predicted proteome. Excluding TCDB class 8.A accessory factors, 680 proteins were classified as canonical transporters and used for downstream functional analyses. Transporter annotation confidence results in 303 high-confidence transporters supported simultaneously by TCDB classification, conserved Pfam domains, and predicted transmembrane topology (Table S10).

Expression analysis (Table S11) confirms that the transporter repertoire is not only numerically expanded but also transcriptionally active, with strong expression of secondary active transporters (TC class 2.A) and ABC transporters (3.A), consistent with active substrate uptake and efflux under osmotrophic conditions.

Functionally, the transporter repertoire encompasses a broad spectrum of nutrient acquisition and homeostatic systems. Proton-coupled Major Facilitator Superfamily transporters (2.A.1; 34 genes) dominate soluble carbohydrate uptake, while amino acid permeases (2.A.3) and peptide transporters support organic nitrogen assimilation. Inorganic nitrogen uptake is enabled by ammonium transporters (1.A.11; AMT/MEP family), indicating direct assimilation of environmental $NH_4^+$. Metal homeostasis is mediated by ZIP ($Zn^{2+}$), NRAMP ($Fe^{2+}$/$Mn^{2+}$), and CTR ($Cu^+$) transporters, complemented by CDF efflux systems for detoxification.

Ion and osmotic regulation are supported by aquaporins, CLC chloride channels, $Na^+$/$H^+$ exchangers, and mechanosensitive Piezo channels, indicating tight control of membrane potential and osmotic stress responses. Primary active transporters (3.A) include ABC transporters, V-type ATPases, and P-type ATPases, providing ATP-driven solute transport and proton motive force generation, while organelle-associated systems (e.g., mitochondrial carriers and protein import complexes) integrate intracellular metabolic exchange.

Notably, 207 transporter-like proteins lack TCDB assignment, representing a major fraction of the predicted transportome and likely reflecting either lineage-specific diversification or highly divergent homologs not captured in current reference databases.

<u>Supplementary Note to results section 6: Phenotypic assays validate broad pentose utilization</u>

Substrate utilization in Biolog Phenotypic MicroArray assays was classified using an approach aligned with the manufacturer's assay design. Substrate utilization was identified by color change, and the quantitative confirmation with endpoint $\Delta OD_{590}$ exceeding the negative control by more than 0.08 (PM1) or 0.05 (PM2) are confirmed by both criteria. Of the 17 positive substrates, 13 were confirmed by both colorimetric and quantitative criteria. Three substrates (dihydroxyacetone, sorbic acid, 2-deoxy-D-ribose) showed visual color development but negative endpoint $\Delta OD_{590}$ due to cell sedimentation. One substrate (L-Rhamnose, PM2; $\Delta OD_{590}$ = 0.075) showed color change but had an endpoint $\Delta OD_{590}$ below the negative control value (0.081), and was classified as positive on the basis of the colorimetric readout alone.

Several abundant carbohydrates produced no colorimetric response: D-glucose ($\Delta OD_{590}$ = 0.026), D-fructose (0.025), D-galactose (0.036), D-mannose (0.034), sucrose, maltose, lactose, and D-cellobiose, all at or below the negative control (PM2 $\Delta OD_{590}$ = 0.081). The absence of a detectable response to free monomeric glucose is unexpected given that *S. psychra* encodes a full glycolytic gene complement and functional MFS transporters. One explanation lies in transporter kinetics: if *S. psychra*'s MFS transporters are calibrated for trace-level dissolved monomers in the dilute organic milieu of lichen substrates, supra-physiological glucose concentrations could paradoxically suppress growth through osmotic effects or catabolite repression of the extracellular enzyme machinery on which this organism depends for nutrient acquisition, a phenomenon well documented in yeasts (Gancedo, 1998). Alternatively, *S. psychra* may preferentially acquire hexoses in oligomeric or polymeric form (e.g. as galactomannan hydrolysate) rather than as free monomers, consistent with the robust utilization of galacto-oligosaccharide derivatives (3-O-β-D-Galactopyranosyl-D-Arabinose, Palatinose) in these assays despite the absence of a free galactose response.

Canonical arabinose isomerase (K01804) is absent despite D-arabinose and L-arabinose utilization (Day 11, $\Delta OD_{590}$ = 0.1640; Day 12, $\Delta OD_{590}$ = 0.1871 respectively), suggesting catabolism via an alternative oxidoreductive route (e.g. non-phosphorlyative oxidative pathway) or through non-canonical isomerization. L-lyxose catabolism in eukaryotes is poorly characterized; the broadly expressed xylose isomerase (K01805; 437.9 TPM) may accommodate lyxose through relaxed substrate specificity. These divergences indicate lineage-specific catabolic strategies warranting biochemical characterization.

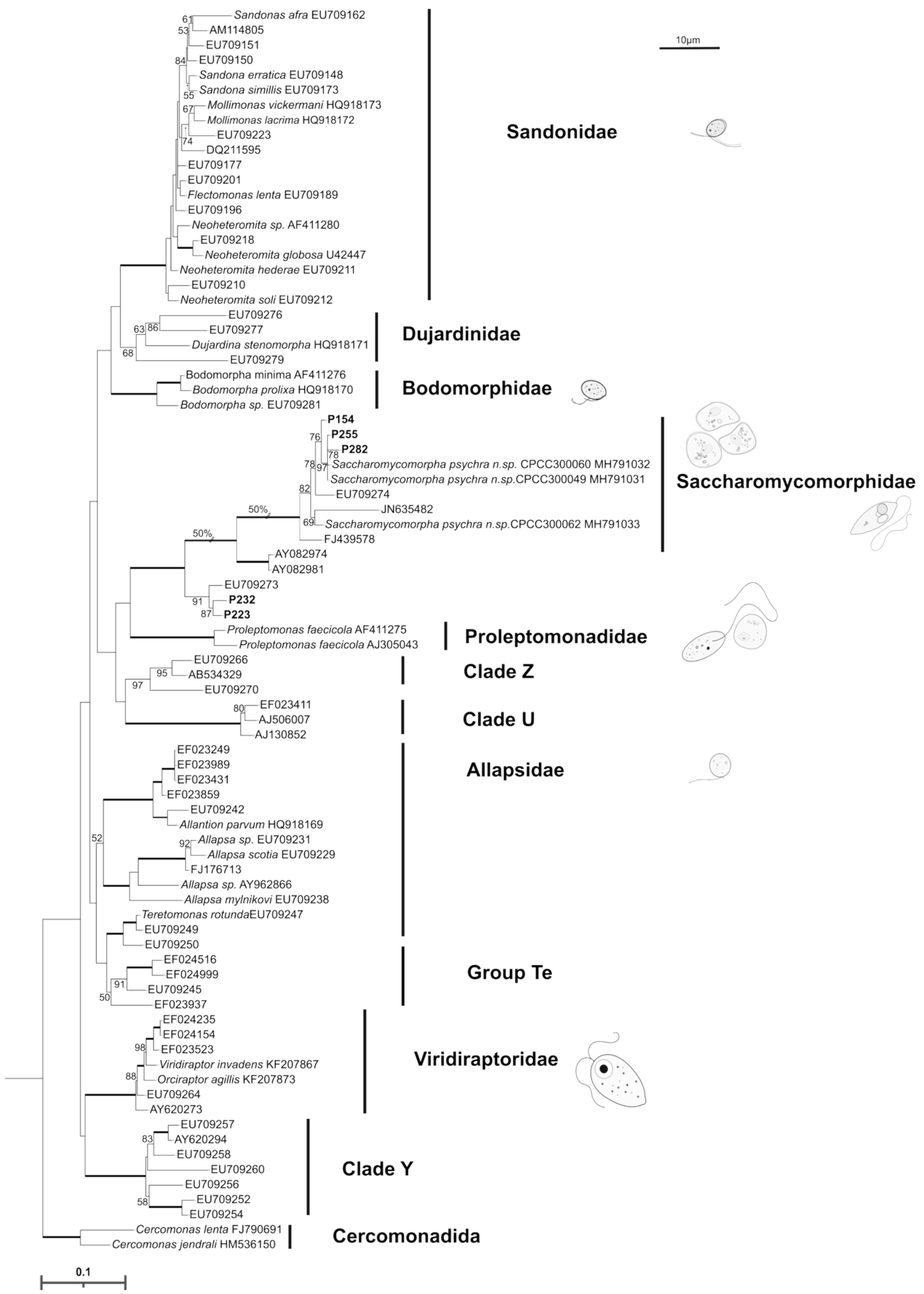


**Figure S1** Phylogenetic placement of *Saccharomycomorpha* isolates based on 18S rRNA gene sequences. Maximum-likelihood phylogeny of representative Glissomonadida inferred from 18S rRNA gene sequences (MAFFT alignment; RAxML, GTR+I+G model; 200 bootstrap replicates). Numbers at nodes indicate bootstrap support (%). Five isolates (P154, P223, P232, P255 and P282) clustered within the family Saccharomycomorphidae. Isolates P255 and P282 grouped with the *Saccharomycomorpha psychra* type strains (MH791031 and MH791032), while P255 showed the highest sequence identity to the type strain (99.65% by BLAST) and was therefore selected for all downstream analyses, including genome sequencing. Sequence is available under SUB16322189; PZ672971.

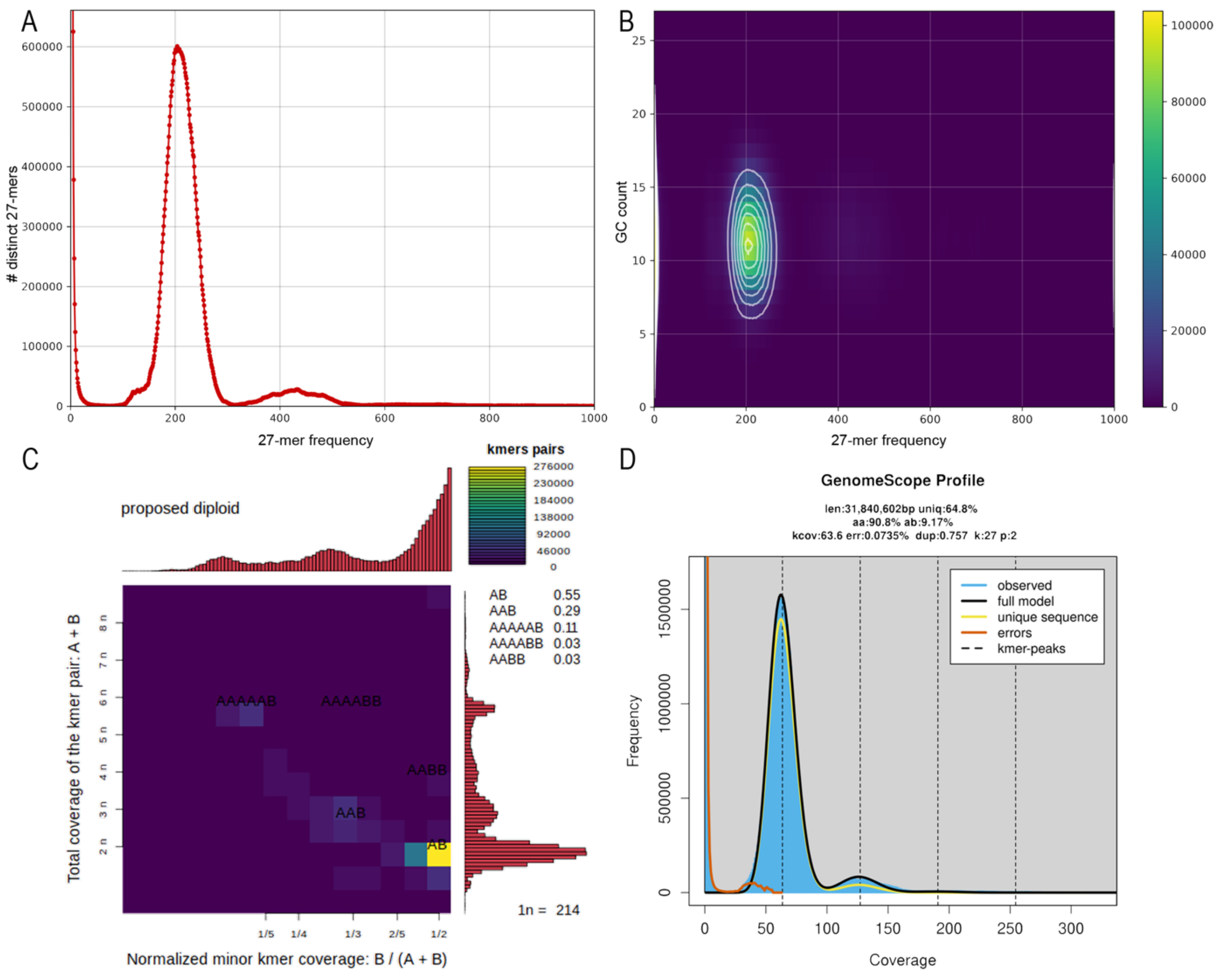


**Figure S2 Genomic property analyses of *Saccharomycomorpha psychra* based on *k*-mers extracted from PacBio HiFi reads (*k*=27).** (A) Two peaks are visible at 205X and 410X corresponding to heterozygous and homozygous *k*-mers, respectively, suggesting a diploid genome, B) The GC content distribution shows no additional high-GC content, suggesting the absence of bacterial contamination in the reads, C) The Smudgeplot shows a distinct smudge of *k*-mers with an A/B configuration, and a smaller region of AA/B configuration, identifying the organism as diploid and indicating duplication, D) Genomescope2.0 analysis indicates diploidy (p=2) with a heterozygosity (ab) of 9.17%. Predicted haploid genome length (len) is ~32 MB. Following Michael C. Schatz (Johns Hopkins University) recommendations (pers. communication) a subset to reduce coverage to approx. 60X was used.

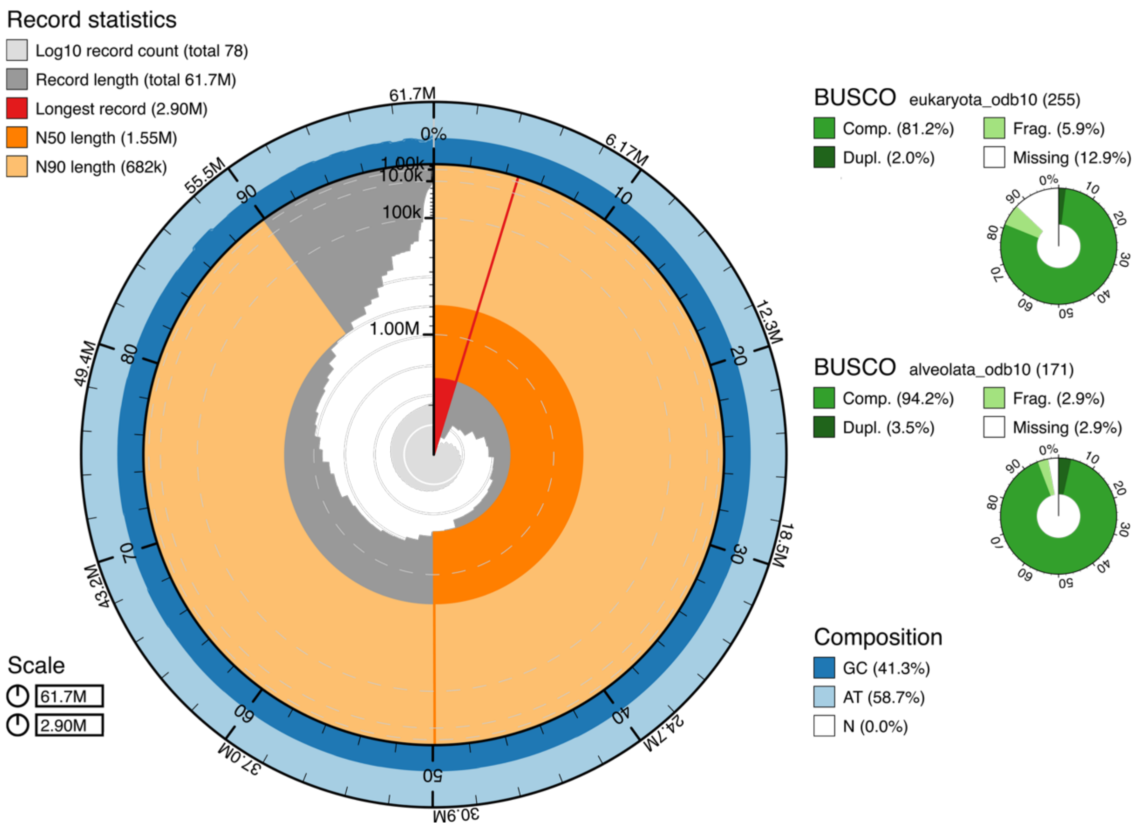


**Figure S3 Snail plot summarizing assembly statistics generated with BlobTools.** The circular plot depicts scaffold length distribution (outer rings), cumulative genome size, GC content distribution, and BUSCO completeness metrics. Most of the sequence contained in relatively large contigs, consistent with the $N_{50}$ of 1.55 Mb. GC content is concentrated around ~40–42%. BUSCO analysis indicates that the majority of expected eukaryotic single-copy orthologs are present, with most identified as complete and single-copy and a small fraction classified as duplicated or fragmented. The BUSCO analysis against the alveolata database presents 94.2% completeness with only 2.9% missing, indicating overall good assembly completeness.

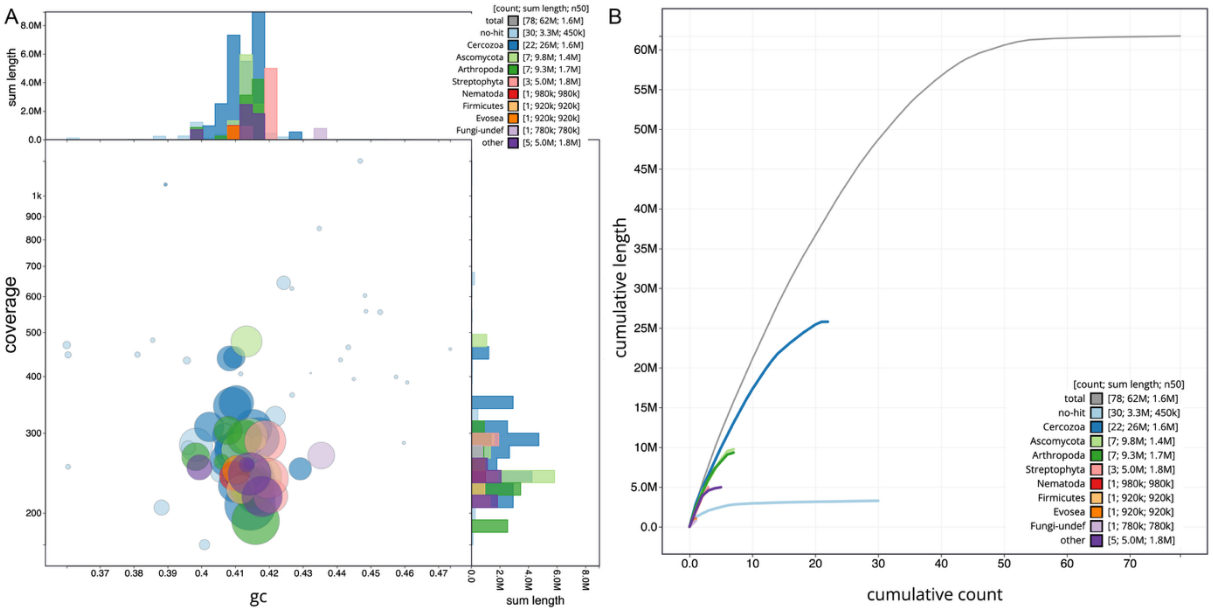


**Figure S4 Genome assembly characteristics inferred using BlobTools.** (A) Blob plot showing GC content versus sequencing coverage for contigs in the *Saccharomycomorpha psychra* genome assembly. Each circle represents a contig, with circle size proportional to contig length and color indicating the best BLAST-based taxonomic assignment. The majority of contigs cluster around ~0.40–0.42 GC and ~200–400X coverage, consistent with the primary genome component. Taxonomic annotation indicates that most of the assembled sequence corresponds to Cercozoa (22 contigs; ~26 Mb), representing the dominant fraction of the assembly. Additional contigs are assigned to Ascomycota (~9.8 Mb across 7 contigs) and Arthropoda (~9.3 Mb across 7 contigs), with smaller contributions from Streptophyta (~5.0 Mb), Nematoda (~0.98 Mb), Firmicutes (~0.92 Mb), Evosea (~0.92 Mb), and unclassified Fungi (~0.78 Mb). These assignments likely reflect biases in reference databases toward model organisms and the limited representation of protistan proteins. A subset of contigs (~3.3 Mb across 30 contigs) lacks significant BLAST hits. The clustering of most contigs within a narrow GC content and coverage range suggests a coherent main genome component. (B) Cumulative scaffold length plotted against cumulative contig count. The assembly comprises 78 scaffolds and 93 contigs with a total length of ~62 Mb and an $N_{50}$ of ~1.6 Mb. The majority of the assembly length derives from Cercozoa-assigned contigs (~26 Mb), followed by smaller cumulative contributions from other taxonomic groups. Contigs without BLAST hits contribute ~3.3 Mb to the assembly.

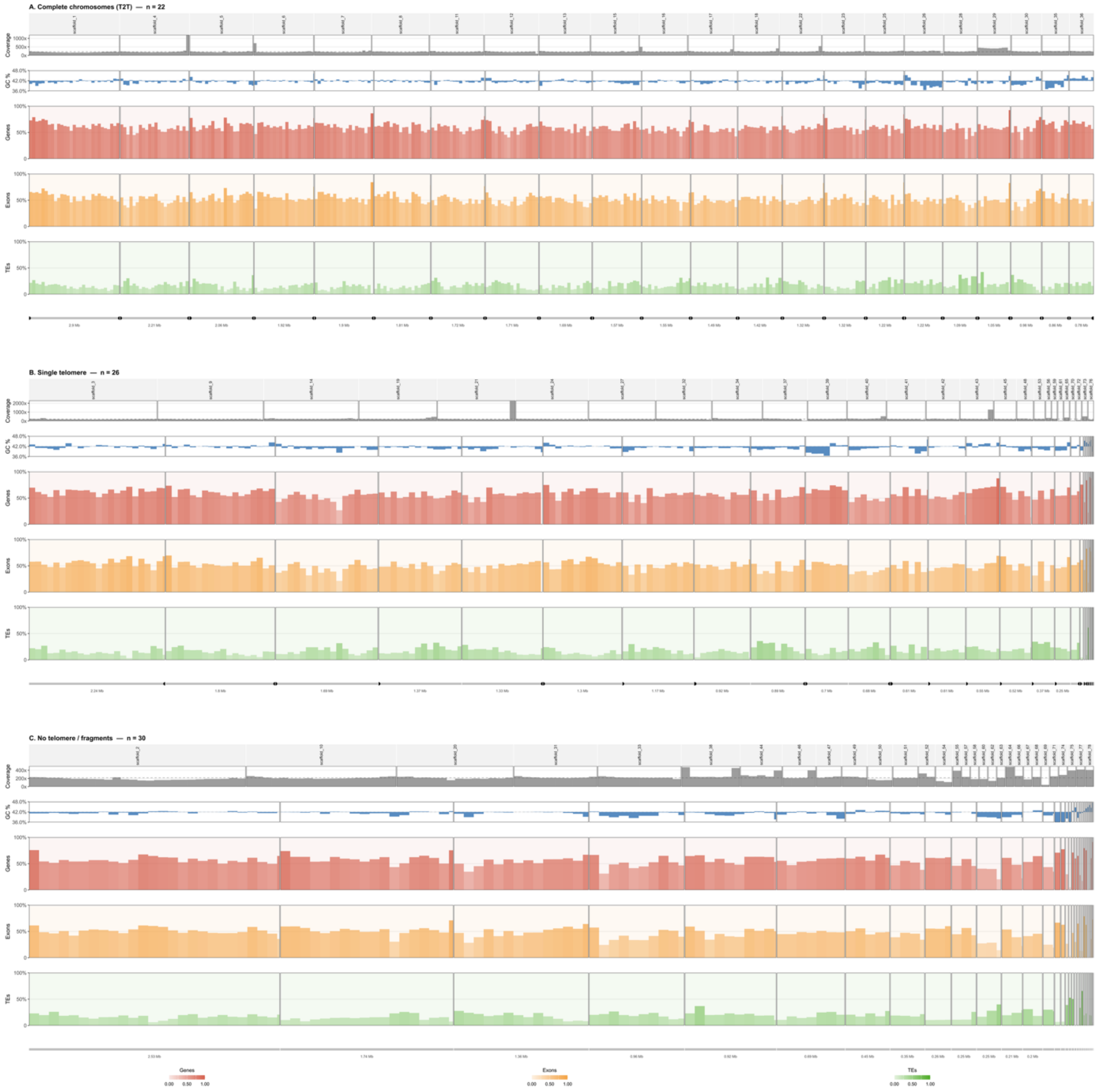


**Figure S5 Genomic properties of *Saccharomycomorpha psychra* assembly.** Genomic landscape tracks (sequencing coverage, GC content, and gene, exon, and transposable element density in 100-kb sliding windows, as in Fig. 1E of the main text) are shown for all 78 assembled scaffolds, grouped into three rows by telomere completeness. Complete chromosomes (n = 22): scaffolds with a telomeric repeat array detected at both termini. Single-telomere scaffolds (n = 26): scaffolds with a telomeric repeat array detected at only one terminus. No-telomere/fragmentary scaffolds (n = 30): scaffolds lacking a detectable telomeric repeat array at either terminus. Track colors follow grey for coverage; blue for GC content; red for gene density; orange for exon density; green for transposable element density.

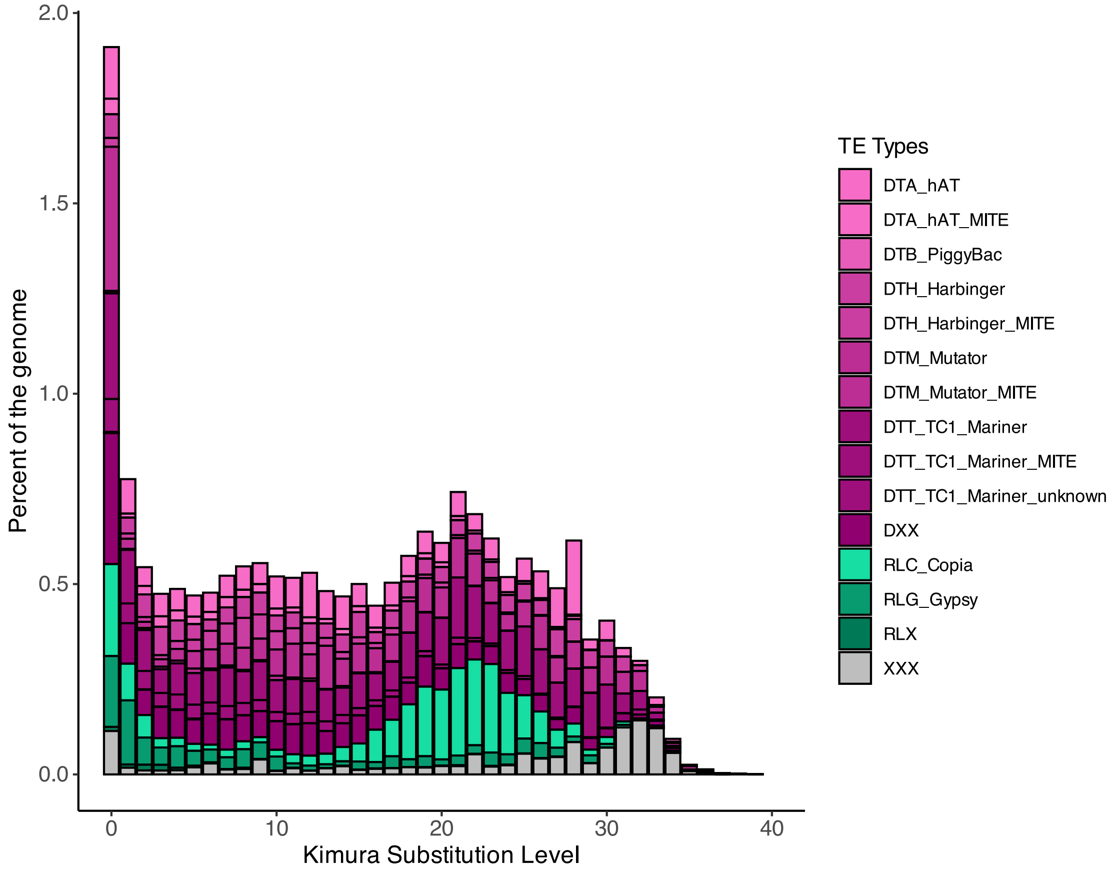


**Figure S6 Transposable element content and Kimura distance-based analysis of historical activity in *Saccharomycomorpha psychra*.** The landscape illustrates the distribution and age-related activity of TE superfamilies, with Kimura substitution level serving as a representative for insertion time. Class I (retrotransposons, green) and Class II (DNA transposons, magenta) elements exhibit fluctuating activity patterns over evolutionary time, reflecting bursts of transposition activity.

**Table S1: Annotation statistics of *Saccharomycomorpha psychra***, before and after deduplication. Deduplication removed redundant transcript isoforms while preserving the complete predicted gene set (17,680 protein-coding genes), yielding a non-redundant annotation used for all downstream functional and expression analyses.

| | **initial** | **deduplicated** |
|---|---|---|
| **Protein-coding genes** | 17680 | 17680 |
| **CDS** | 32406 | 30543 |
| **mRNA** | 18511 | 17680 |
| **Intron** | 13921 | 12889 |
| **Exon** | 32406 | 30543 |
| **Start codon** | 18482 | 17652 |
| **Stop codon** | 18498 | 17667 |
| **functional hits via eggNOG** | | 9968 |
| **Number of secreted proteins** | | 1015 |
| **[%] of proteome** | | 5,75 |

**Table S2: Comparison of publicly available rhizarian genome assemblies.** Assembly statistics and BUSCO completeness values were recalculated where possible using BUSCO v5 with the Eukaryota odb12 lineage dataset (n: 129) to enable standardized comparison across assemblies. Consequently, completeness estimates may differ from those reported in the original publications, including the values reported elsewhere in this manuscript for *Saccharomycomorpha psychra*, which were generated using the Eukaryota odb10 dataset. The genome assembly of *Ammonia veneta* was not publicly available at the time of analysis and therefore retains the BUSCO completeness reported in the original publication.

| **Species** | **Genome size (Mb)** | **Contigs (n)** | **Contig $N_{50}$ (Mb)** | **Protein- coding genes** | **C [%]** | **S [%]** | **D [%]** | **F [%]** | **M [%]** | **Lifestyle** | **Reference** |
|---|---|---|---|---|---|---|---|---|---|---|---|
| ***Saccharomycomorpha psychra*** | 61.7 | 93 | 1.329 | 17,680 | 74.4 | 72.9 | 1.6 | 5.4 | 20.2 | Free-living | this study |
| ***Ammonia veneta*** | 433 | 284,473 | 0.05 | 350,650 | 80.0 | – | – | – | – | Foraminifer | Ishitani et al., 2025 |
| ***Mikrocytos mackini*** | 49.7 | 16,029 | 0.005 | 14,372 | 10.6 | 8.6 | 2.0 | 5.9 | 83.5 | Pathogen/parasite | Zarsky et al., 2023 |
| ***Paramikrocytos canceri*** | 12.7 | 3,113 | 0.007 | 8,201 | 5.9 | 5.9 | 0.0 | 4.3 | 89.8 | Pathogen/parasite | Onuț-Brännström et al., 2023 |
| ***Paulinella micropora*** | 707.1 | 7,048 | 0.14 | 32,361 | 18.4 | 17.3 | 1.2 | 19.6 | 62.0 | Free-living testate amoeba | Lhee et al., 2021 |
| ***Plasmodiophora brassicae*** | 24.8 | 1,172 | 0.510 | 9,913 | 84.7 | 80.0 | 4.7 | 7.1 | 8.2 | Pathogen/parasite | Schwelm et al., 2015 |
| ***Reticulomyxa filosa*** | 101.9 | 50,809 | 0.004 | 40,160 | 42.4 | 38.8 | 3.5 | 27.8 | 29.8 | Foraminifer | Glöckner et al., 2014 |
| ***Lotharella oceanica*** | 0.68 | 5 | 0.194 | 668 | 0.8 | 0.8 | 0.0 | 1.2 | 98.0 | Chlorarachniophyte (alga) | Tanifuji et al., 2014 |
| ***Bigelowiella natans*** | 91.4 | 3,736 | 0.059 | 22,320 | 38.0 | 36.9 | 1.2 | 22.0 | 40.0 | Chlorarachniophyte (alga) | Curtis et al., 2012 |
| ***Marteilia pararefringens*** | 22.35 | 4,636 | 0.006 | 5,208 | 10.1 | 10.1 | 0 | 21.7 | 68.2 | Parasite (Paramyxida, bivalve) | Hiltunen Thorén et al., 2024 |
| ***Bonamia ostreae*** | 10.29 | 7,958 | 0.001 | 9,694 | 17.8 | 17.8 | 0 | 17.8 | 64.3 | Parasite (Haplosporida, bivalve) | Hiltunen Thorén et al., 2024 |
| ***Paramarteilia canceri*** | 36.81 | 16,785 | 0.003 | 2,340 | 10.9 | 10.9 | 0 | 24.8 | 64.3 | Parasite (Paramyxida, crustacean) | Hiltunen Thorén et al., 2024 |
| ***Cercozoa sp.* M6MM** | 27.0 | 15 | 5.83 | 9,694 | 11.6 | 11.6 | 0 | 31.8 | 56.6 | Amphizoic amoeba | Hiltunen Thorén et al., 2024 |

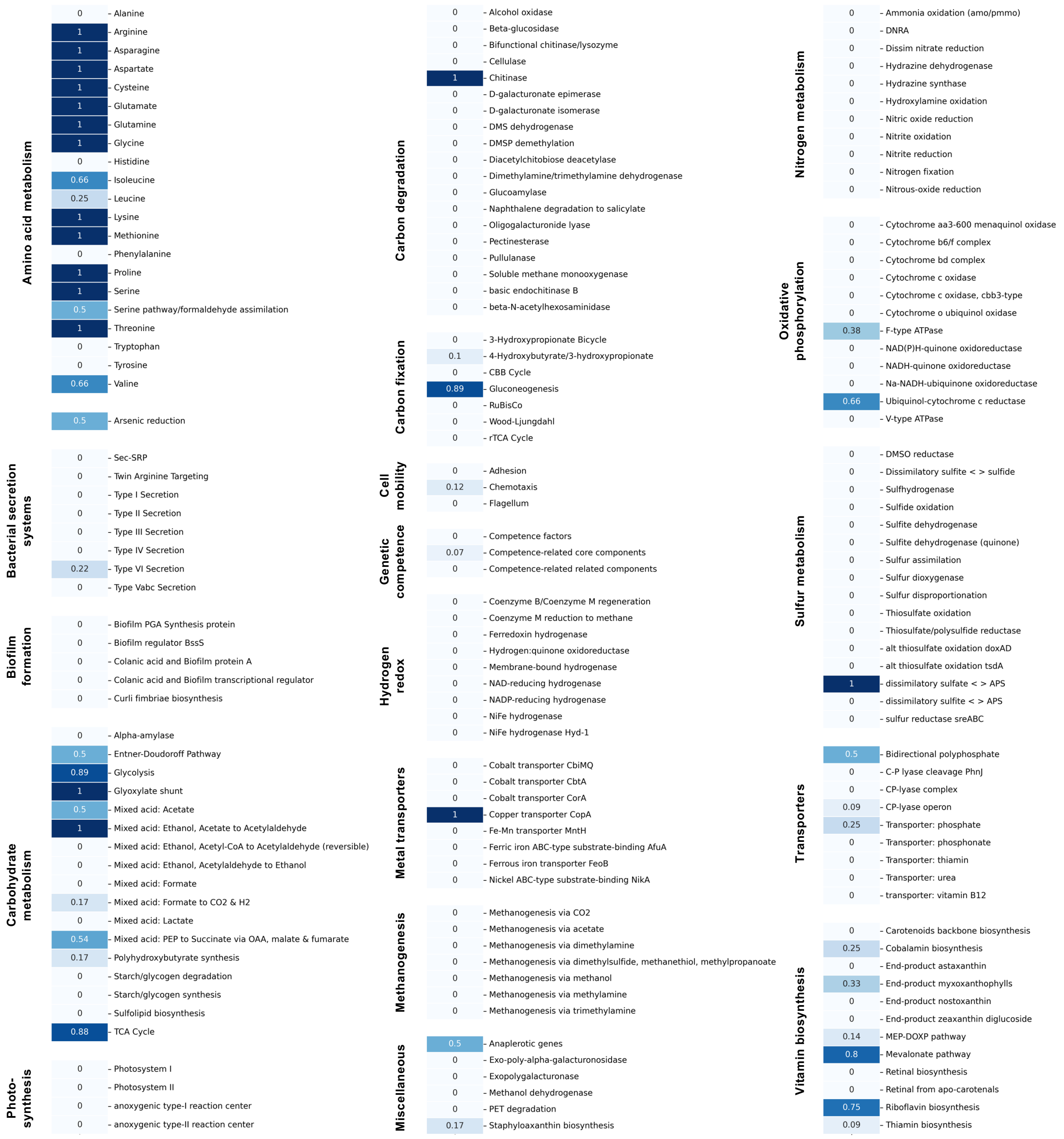


**Figure S7: Heatmap of KEGG pathway completeness in *Saccharomycomorpha psychra*.** Metabolic pathways reconstructed from KEGG Orthology assignments are grouped by functional category. Pathway completeness (0–1) is indicated by blue color intensity (lighter = lower completeness, darker = higher completeness). The reconstruction highlights extensive coverage of central carbon metabolism, amino acid biosynthesis, and energy metabolism.

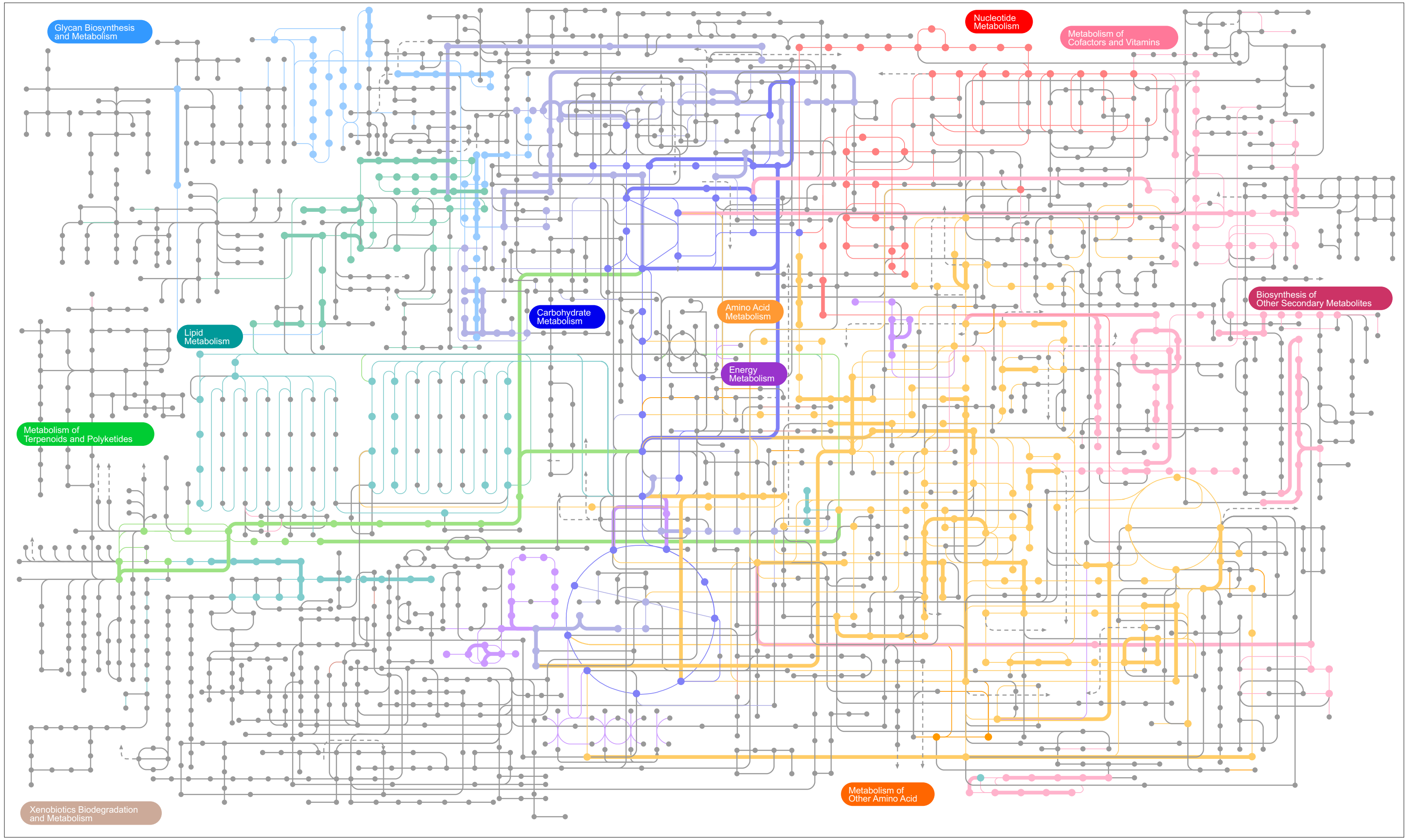


**Figure S8 Metabolic pathway map of *Saccharomycomorpha psychra*** generated with iPath3.0. KEGG Orthology (KO) assignments were obtained using KofamKOALA for functional annotation of protein sequences, with transcript abundance estimated via Salmon. Colored metabolic category and sized by expression level: thin lines represent presence of KO terms, thick lines indicate ≥1 TPM (expressed genes). An interactive version of this reconstruction can be accessed via https://github.com/HuesnaOeztoprak/Saccharomycomorpha/tree/main/Metabolism.

**Table S3: Top20-expressed secreted proteins of *Saccharomycomorpha psychra*.** Secreted proteases dominate the most highly expressed proteins, together with several abundant proteins lacking functional annotation, highlighting both active extracellular protein degradation and the presence of potentially novel secreted factors.

| Function | Gene | TPM | logTPM |
|---|---|---|---|
| Pro-kumamolisin, activation domain | g10993 | 8250.1897 | 3.9165 |
| Cysteine-type peptidase activity | g9335 | 6816.6665 | 3.8336 |
| Not annotated | g10160 | 5294.9888 | 3.7239 |
| Insulin-like growth factor 2 receptor | g10482 | 4103.6554 | 3.6133 |
| Cysteine-type endopeptidase activity | g12933 | 3713.5851 | 3.5699 |
| acr, cog1565 (uncharacterized) | g4560 | 3465.6610 | 3.5399 |
| Peptidase S10 family | g10102 | 3239.9595 | 3.5107 |
| Not annotated | g10612 | 3207.8444 | 3.5063 |
| Cell wall organization | g4400 | 3028.8988 | 3.4814 |
| Not annotated | g4382 | 2695.8647 | 3.4309 |
| Peptidase C1 family | g7039 | 2574.3755 | 3.4108 |
| Carboxylesterase/lipase | g12719 | 2540.0083 | 3.4050 |
| Not annotated | g12056 | 2331.6034 | 3.3678 |
| Not annotated | g2567 | 2297.5824 | 3.3615 |
| Not annotated | g9787 | 1751.7786 | 3.2437 |
| Unfolded protein binding (Calreticulin) | g11482 | 1742.4585 | 3.2414 |
| acr, cog1565 (uncharacterized) | g4498 | 1735.7053 | 3.2397 |
| Cysteine-type peptidase activity | g16204 | 1580.6731 | 3.1991 |
| Pro-kumamolisin, activation domain | g15452 | 1487.3105 | 3.1726 |
| Not annotated | g16218 | 1460.9554 | 3.1649 |

**Table S4. Genome-wide expression summary of functional groups involved in secretion in *Saccharomycomorpha psychra*.** Secreted proteins, transporters and CAZymes together account for a disproportionate fraction of transcript abundance relative to their genomic representation, consistent with an actively osmotrophic lifestyle. Note: varying number of genes i.e. secreted CAZymes (60 of 64 annotated genes) or gene total reflects genes recovered in the Salmon quantification matrix (17,066 of 17,680 annotated genes); 614 genes lacking expression are excluded from expression analyses.

| Category | Genes (n) | % of genes | Sum TPM | % of TPM | Mean TPM | Median TPM | % zero-expressed |
|---|---|---|---|---|---|---|---|
| background | 15,493 | 90.80% | 843,233 | 84.70% | 54.4 | 5.0 | 18.2% |
| Secreted | 908 | 5.32% | 113,925 | 11.44% | 126.0 | 2.4 | 15.6% |
| Transporter | 568 | 3.33% | 33,286 | 3.34% | 58.6 | 20.3 | 1.9% |
| Secreted CAZyme | 60 | 0.35% | 3,769 | 0.38% | 62.8 | 33.0 | 5.0% |
| Non-secreted CAZyme | 37 | 0.22% | 1,337 | 0.13% | 36.1 | 13.4 | 0% |
| TOTAL | 17,066 | 100% | 995,549 | 100% | — | — | — |

**Table S5. Pairwise Wilcoxon rank-sum tests with effect sizes (BH-corrected) of functional groups of *Saccharomycomorpha psychra*.** Secreted CAZymes and transporters show consistently elevated expression relative to the genomic background, whereas effect sizes remain modest because of the large and heterogeneous background gene set (the Fisher's exact test odds ratios (CAZyme OR=3.15, Secreted OR=2.11) provide complementary evidence of biologically meaningful enrichment). Note: varying number of genes are due to the difference of genomic prediction and expression.

| Group 1 | Group 2 | n1 | n2 | p.adj | Significance | r (effect size) |
|---|---|---|---|---|---|---|
| background | CAZyme | 15,493 | 37 | 0.0020 | ** | 0.027 |
| background | Secreted | 15,493 | 908 | 0.864 | ns | 0.001 |
| background | Secreted CAZyme | 15,493 | 60 | $1.2 \times 10^{-9}$ | **** | 0.050 |
| background | Transporter | 15,493 | 568 | $2.7 \times 10^{-59}$ | **** | 0.129 |
| CAZyme | Secreted | 37 | 908 | 0.002 | ** | 0.108 |
| CAZyme | Secreted CAZyme | 37 | 60 | 0.034 | * | 0.230 |
| CAZyme | Transporter | 37 | 568 | 0.127 | ns | 0.064 |
| Secreted | Secreted CAZyme | 908 | 60 | $3.95 \times 10^{-7}$ | **** | 0.169 |
| Secreted | Transporter | 908 | 568 | $1.52 \times 10^{-32}$ | **** | 0.313 |
| Secreted CAZyme | Transporter | 60 | 568 | 0.070 | ns | 0.076 |

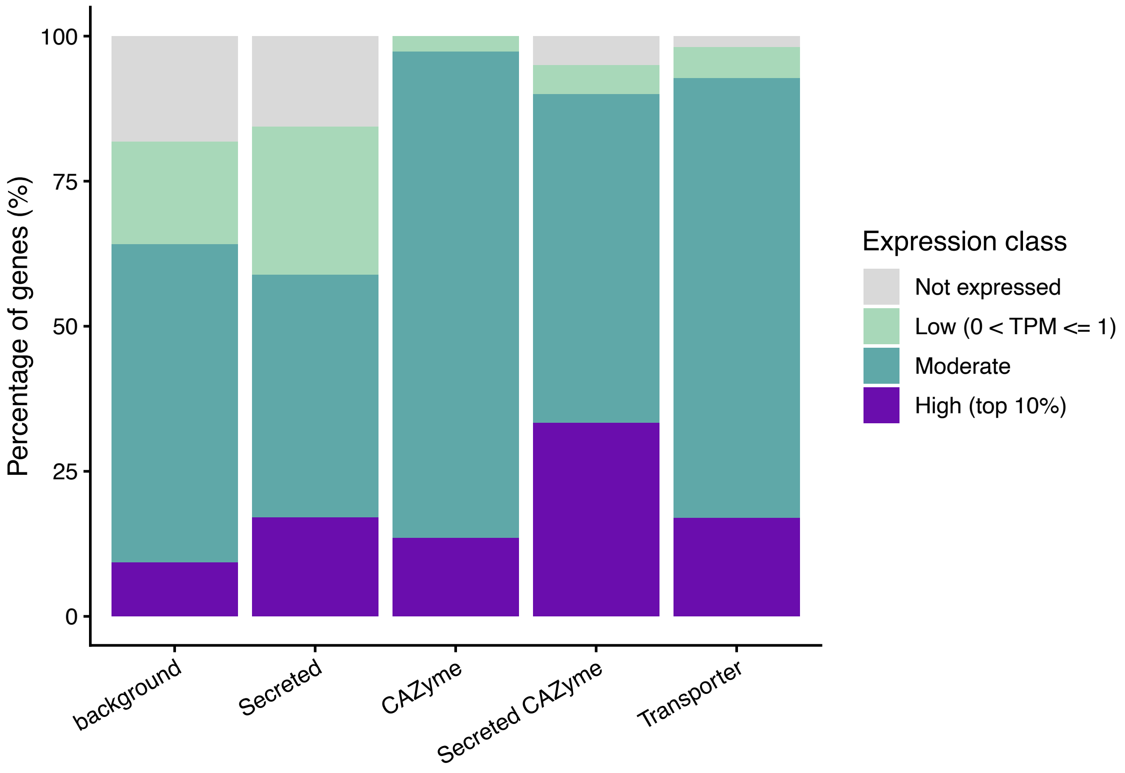


**Figure S9 Expression breadth by functional category**. Percentage of genes in each category falling into four expression tiers: not expressed (TPM = 0), low (0 < TPM ≤ 1), moderate (1 < TPM ≤ 69.4), high (top 10%, TPM > 69.4). Key result: the 'Secreted' category has a distinctly different profile from all other categories: the highest proportion of unexpressed genes (14.9%) combined with the highest proportion in the top-10% tier (18.1%), confirming the bimodal distribution seen in Panel B of Fig.3 (main text). This pattern suggests the secretome contains both constitutively active enzymes and a large reservoir of condition-specific or developmental effectors not expressed under the sampled culture conditions.

**Table S6: Methodical filtering and identification of CAZymes** with Phobius, SignalP, DeepLoc and dbCAN2. Combining dbCAN2 predictions with secretion and localization filters reduced 916 initial candidate annotations to 109 high-confidence CAZyme genes, providing a conservative dataset for downstream analyses.

| | Number of genes |
|---|---|
| **Hotpep.out** | 249 |
| **diamond.out** | 624 |
| **hmmer.out** | 220 |
| **signalp.out** | 2250 |
| **Total hits** | 916 |
| **high confidence (more than 2 tools)** | 109 |

**Table S7: Representative activity and function of CAZymes of *Saccharomycomorpha psychra* and their ecological implications.** The repertoire is dominated by enzymes targeting fungal cell walls, galactomannans and plant-derived polysaccharides, supporting a broad capacity for extracellular degradation within lichen thalli.

| CAZyme family | Representative activity | Function | Putative ecological implication |
|---|---|---|---|
| GH3 | β-glucosidases, β-xylosidases, N-acetyl-β-glucosaminidases | Hemicellulose degradation; oligosaccharide hydrolysis; arabinogalactan turnover | Releases glucose from cellobiose and β-glucans during degradation of plant-derived organic matter |
| GH5_9 | Endoglucanases, β-mannanases | Cellulose and β-mannan degradation | Degradation of cellulose-rich plant material and some fungal β-glucans |
| GH16 | β-1,3/1,4-glucanases, lichenases | Mixed-linkage glucan hydrolysis | Degradation of lichenan and other β-glucans from lichen thalli and microbial cell walls |
| GH27 | α-galactosidase | Galactomannan side-chain removal | Hydrolysis of galactose residues from plant galacto(gluco)mannans |
| GH29 | α-L-fucosidase | Fucose removal | Degradation of fucosylated glycans from plants and microbial glycoproteins |
| GH31 | α-glucosidases, α-xylosidases | α-linked glucan and xyloglucan hydrolysis | Degradation of xyloglucan and terminal α-glucosides |
| GH35 | β-galactosidase | Galactan degradation | Breakdown of galactans and arabinogalactans from plant cell walls |
| GH43_24 | Arabinofuranosidases, xylosidases | Arabinoxylan degradation | Hydrolysis of arabinose-rich hemicelluloses |
| GH51 | α-L-arabinofuranosidase | Arabinan degradation | Removal of arabinose side chains from hemicellulose and pectin |
| GH78 (+CBM67) | α-L-rhamnosidase | Rhamnogalacturonan degradation | Cleavage of rhamnose-containing pectic polysaccharides |
| GH88 | Unsaturated glucuronyl hydrolase | Pectin catabolism | Utilization of pectin degradation products |
| GH141 | α-L-fucosidase | Fucose metabolism | Turnover of fucosylated glycans |
| CE15 | Glucuronoyl esterase | Lignin-carbohydrate complex cleavage | Releases hemicellulose from lignin during lignocellulose degradation |
| GH38 | α-mannosidase | Mannose trimming | Processing of N-glycans and fungal galactomannans |
| GH47 | α-1,2-mannosidase | Early N-glycan maturation | Glycoprotein processing and mannan degradation |
| GH125 | Exo-α-mannosidase | Terminal mannose removal | Mannan utilization and glycoprotein turnover |
| GH20 | β-N-acetylhexosaminidase | Terminal GlcNAc removal | Glycoprotein degradation and chitin recycling |
| GH18 | Chitinase | Chitin hydrolysis | Digestion of fungal cell walls, including lichen mycobionts |
| GH89 | α-N-acetylglucosaminidase | Aminosugar metabolism | Glycan remodeling and glycoprotein turnover |
| GH31 (subset) | α-glucosidase | Glycan trimming | Processing of N-linked glycans |
| GT2 | Cellulose synthase, chitin synthase | Structural polysaccharide synthesis | Cell-wall/extracellular matrix biosynthesis |
| GT4 | Glucosyltransferases | Glycan elongation | Cell-surface glycoconjugate synthesis |

| | | | |
|---|---|---|---|
| GT13 | β-1,4-galactosyltransferase | Galactosylation | Protein glycosylation |
| GT20 | Trehalose-6-phosphate synthase | Trehalose biosynthesis | Osmoprotection and cold/desiccation tolerance |
| GT22 | Dolichyl-phosphate-mannose synthase | Dolichol-linked glycosylation | N-glycan precursor synthesis |
| GT24 | Glucosylceramide synthase | Membrane glycolipid synthesis | Cell membrane remodeling |
| GT41 | O-GlcNAc transferase | Intracellular glycosylation | Regulation of protein activity |
| GT48 | β-1,3-glucan synthase | β-glucan synthesis | Extracellular matrix/cell-wall production |
| GT57 | α-1,3-glucosyltransferase | Glycan assembly | Cell-surface polysaccharide biosynthesis |
| GT66 | Oligosaccharyltransferase | N-glycosylation | Protein maturation |
| GT77 | α-xylosyltransferase | Xylosylation | Glycoprotein and polysaccharide synthesis |
| GT84 | β-glucuronyltransferase | Glucuronylation | Glycoconjugate formation |
| GT58 | Dolichol-P-Man-dependent mannosyltransferase | Mannosyl transfer | Protein glycosylation |
| AA2 | Class II peroxidase | Oxidative lignin modification | Oxidative degradation of lignified plant material |
| AA3_2 | GMC oxidoreductases | Sugar oxidation | Oxidative carbohydrate metabolism and extracellular redox cycling |
| AA4 | Vanillyl alcohol oxidase | Phenolic oxidation | Lignin-derived aromatic compound metabolism |
| AA7 | Glucooligosaccharide oxidase | Oligosaccharide oxidation | Oxidative processing of soluble carbohydrates |
| CBM13 | Xylan/chitin-binding | Glycan targeting | Enhances enzyme binding to plant and fungal polysaccharides |
| CBM35 | Xylan/galactose-binding | Substrate targeting | Improves degradation of hemicellulose |
| CBM51 | Galactose-binding | Glycan recognition | Recognition of galactose-rich glycans |
| CBM57 | β-1,3-glucan-binding | β-glucan targeting | Enhances fungal β-glucan degradation |
| CBM67 | L-rhamnose-binding | Rhamnose targeting | Targets pectic rhamnogalacturonan |
| CBM48 | Glycogen/starch-binding | α-glucan binding | Utilization of storage polysaccharides |
| CBM18 | Chitin-binding | Chitin recognition | Facilitates chitin degradation |
| CBM20 | Starch-binding | Starch binding | Utilization of starch reserves |

**Table S8 *Saccharomycomorpha psychra*'s CAZymes repertoire categorized in substrate-oriented modules.** Grouping CAZymes by substrate reveals that glycoprotein/galactomannan processing and hemicellulose degradation together account for nearly 70% of total CAZyme transcription, consistent with fungal biomass representing a major nutritional resource. Note: The two substrate modules listed here together correspond to the single 'fungal cell wall and glycoprotein' module described in the main text (results); they are separated here to resolve the distinct chitin/β-glucan-targeting and mannan/glycoprotein-targeting gene sets.

| Substrate module | Key families | Total TPM | % of all CAZyme TPM |
|---|---|---|---|
| **Glycoprotein/galactomannan processing** | GH38, GH31, GH20, GH27, GH29, GH47, GH125 | 2,372 | 41.5% |
| **Hemicellulose/pectin/lichen** | GH35, GH3, GH51, GH31, GH43, GH16, CBM67 | 1,592 | 27.9% |
| **Oxidative module** | AA2, AA4, AA3_2, CE15 | 509 | 8.9% |
| **Fungal cell wall β-glucan** | GH5_9, GH18 | 369 | 6.5% |
| **Biosynthetic GTs** | GT20, GT66, GT48, etc. | ~395 | 6.9% |
| **Other** | starch, trehalose, misc | ~471 | 8.3% |

**Table S9: Top20-expressed CAZymes of *Saccharomycomorpha psychra*.** Depicted are gene-ids, CAZyme families and transcripts per million (TPM). Highly expressed enzymes are dominated by GH38 α-mannosidases and secreted glycan-processing enzymes, supporting active degradation of fungal glycoproteins and galactomannan-rich substrates.

| **Gene** | **Family** | **Secreted** | **TPM** | **logTPM** |
|---|---|---|---|---|
| **g2583** | AA2 | FALSE | 371.9941 | 2.5717 |
| **g11077** | GH38 | TRUE | 314.7982 | 2.4994 |
| **g11078** | GH38 | TRUE | 265.5552 | 2.4258 |
| **g547** | GH38 | TRUE | 247.7394 | 2.3957 |
| **g9969** | GH35 | FALSE | 243.4216 | 2.3881 |
| **g7663** | GH13_8 | FALSE | 224.4756 | 2.3531 |
| **g6534** | GH38 | TRUE | 196.0354 | 2.2945 |
| **g13409** | CBM67 | TRUE | 186.4723 | 2.2729 |
| **g12996** | GH20 | TRUE | 184.2877 | 2.2678 |
| **g8016** | GH3 | TRUE | 159.3621 | 2.2051 |
| **g5081** | GT20 | FALSE | 135.5853 | 2.1354 |
| **g1243** | GT66 | FALSE | 130.0006 | 2.1173 |
| **g12444** | GH29 | TRUE | 128.6035 | 2.1126 |
| **g5939** | GH15 | FALSE | 121.8318 | 2.0893 |
| **g6990** | GH5_9 | FALSE | 117.1335 | 2.0724 |
| **g16067** | GH3 | TRUE | 115.4166 | 2.0660 |
| **g7351** | GH51 | TRUE | 115.2499 | 2.0654 |
| **g4083** | AA4 | FALSE | 115.0027 | 2.0645 |
| **g17135** | GH31 | TRUE | 110.6926 | 2.0480 |
| **g2886** | GH31 | TRUE | 108.5950 | 2.0398 |

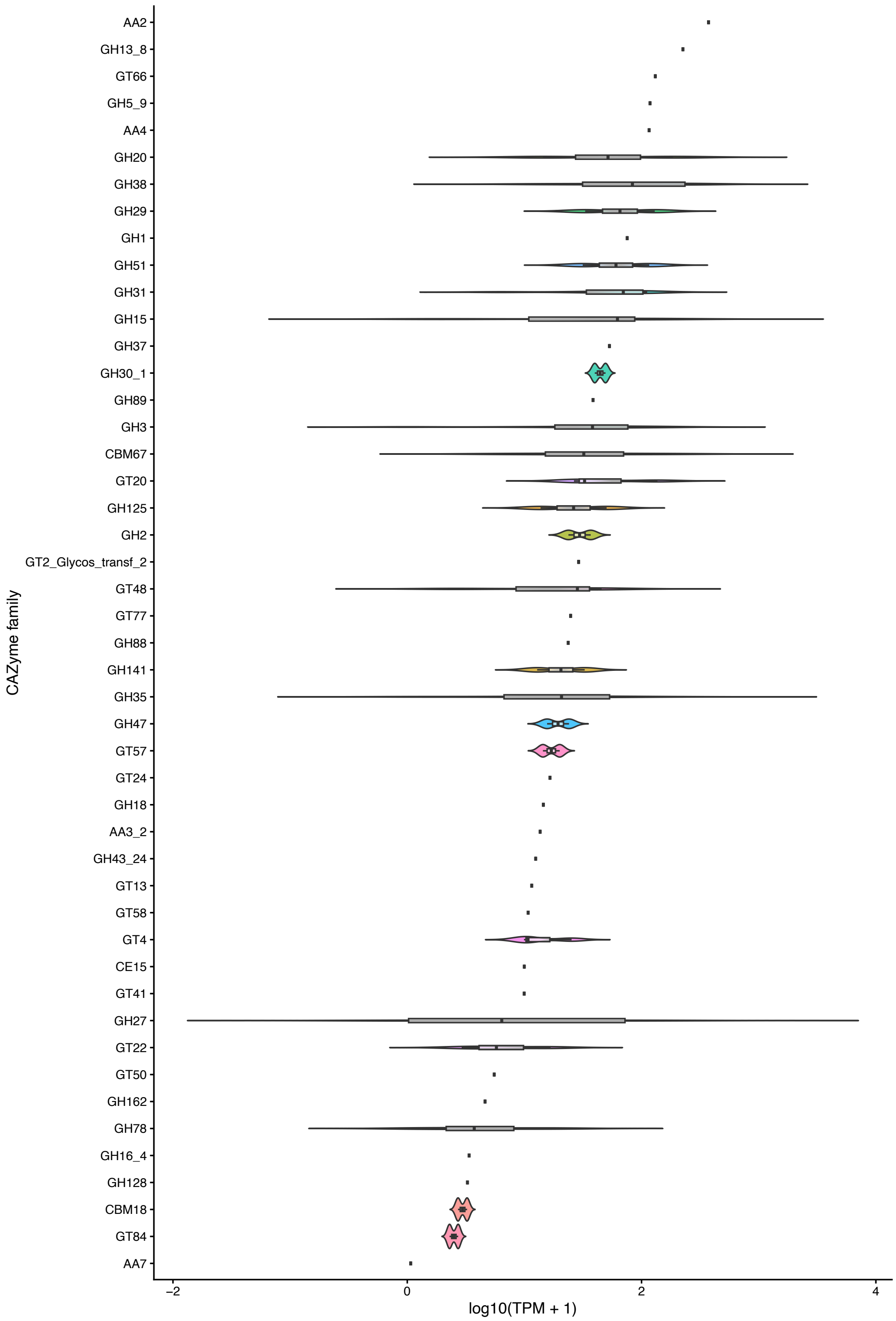


**Figure S10 Per-family expression distribution per CAZyme family.** Violin plots show the distribution of log10(TPM + 1) values across genes within each of the 47 CAZyme families, with overlaid boxplots summarizing the median and interquartile range. Families are ordered by median expression. Because many families contain only one or two genes, several distributions collapse to single observations or very narrow ranges.

**Table S10: Methodical filtering and prediction of Transporterproteins.** High confidence = supported by all three lines of evidence (TCDB hit, Pfam domain, predicted transmembrane topology); medium = two lines of evidence; low = TM helix prediction only.

| | |
|---|---|
| **number of proteins** | 724 |
| **number of proteins excluding TC 8.A accessory factors** | 680 |
| **of total proteome [%]** | 3,85 |
| **high confidence** | 303 |
| **including medium confidence** | 362 |
| **including low confidence** | 724 |

**Table S11: Transporter Classification Database (TCDB) categories identified in the *Saccharomycomorpha psychra* genome.** Secondary active transporters, predominantly members of the major facilitator superfamily (MFS), constitute the largest annotated transporter class, consistent with proton-coupled uptake of extracellular digestion products characteristic of osmotrophic metabolism.

| TCDB class | Functional category | n genes | total TPM | mean TPM | Representative function |
|---|---|---|---|---|---|
| 2.A | Secondary active transporters (MFS, AA permeases, PTR, etc.) | 112 | 7385.7 | 65.9 | Uptake of sugars, amino acids, peptides, ions |
| 3.A | Primary active ABC transporters | 66 | 4109.3 | 62.3 | Lipid, drug, metabolite transport |
| 1.A | Ion channels | 40 | 2147.3 | 53.7 | Ion flux, membrane potential regulation |
| 9.A | Regulatory membrane proteins | 34 | 872.9 | 25.7 | Signaling and transport modulation |
| 9.B | Regulatory membrane proteins | 10 | 292.0 | 29.2 | Signaling complexes |
| 1.B | Porins | 3 | 1000.3 | 333.4 | Large solute transport |
| 3.D | Protein translocases | 3 | 221.8 | 73.9 | Protein secretion/import |
| 4.C | Transport electron carriers | 1 | 208.2 | 208.2 | Electron transfer across membranes |
| 8.A | Auxiliary transport proteins | 1 | 29.3 | 29.3 | Transport modulation |
| 5.B | Uncharacterized transporters | 4 | 21.9 | 5.5 | Putative transport-related proteins |
| 1.I | Mechanosensitive channels | 1 | 48.4 | 48.4 | Osmotic stress response |
| Unknown | Unclassified membrane proteins | 207 | 10611.9 | 51.3 | Putative novel transport systems |

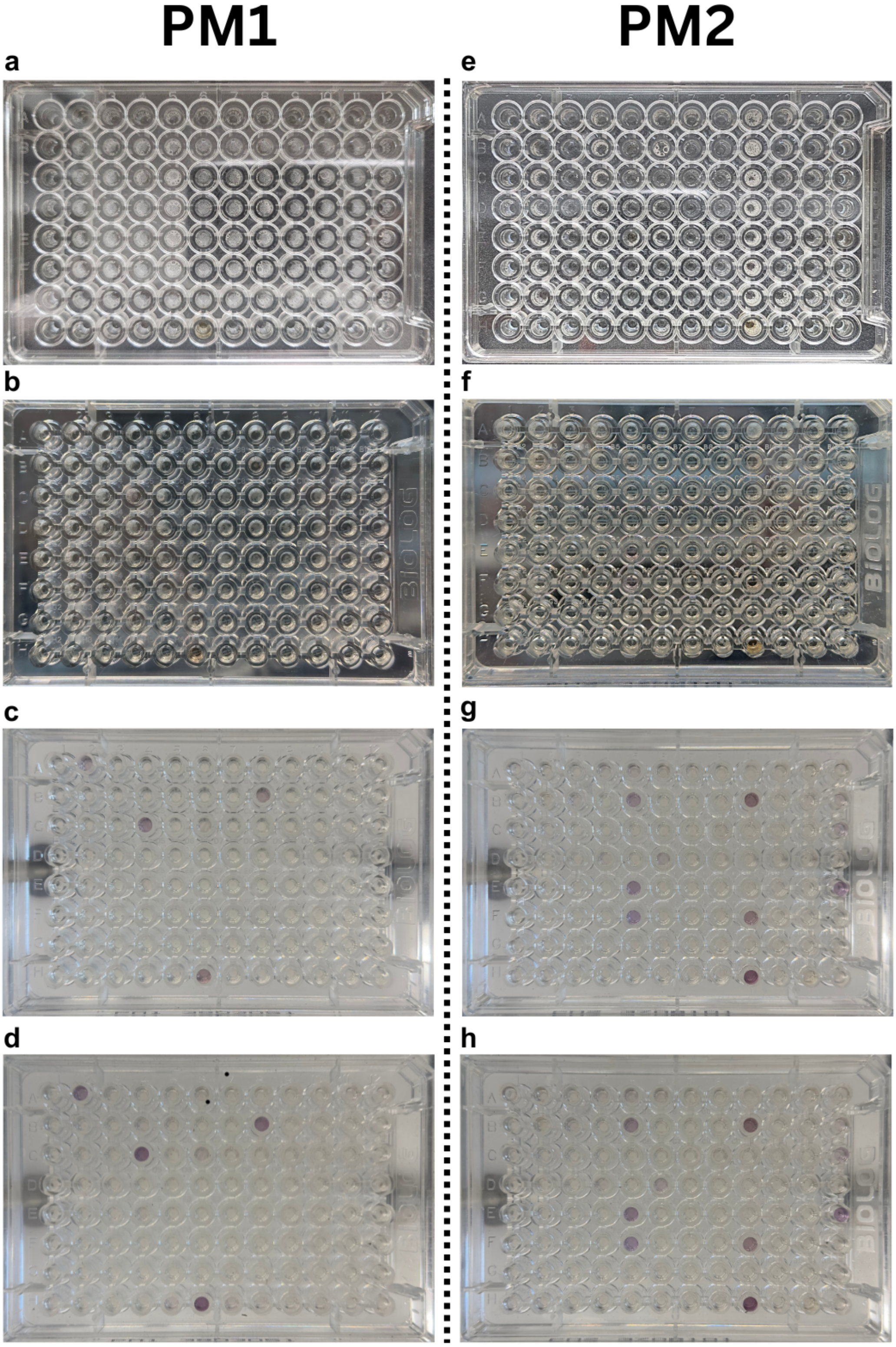


**Figure S11 Time course of color development on Biolog Phenotype MicroArrays$^{TM}$ PM1 (left) and PM2 (right) carbon utilization assays inoculated with *Saccharomycomorpha psychra* and incubated at 8 °C.** Panes (a-d) show PM1 at Day 1 (initial state), Day 4 (first visible color change), Day 17 (last visible color change), and Day 67 (final measurement); while panes (e-h) show PM2 at Day 1 (initial state) ), Day 4 (first visible color change), Day 63 (last visible color change), and Day 67 (final measurement). Color development reflects metabolic activity through reduction of the tetrazolium redox dye, indicating active utilization of the corresponding carbon source in that well.

**Table S12 Biolog Phenotypic MicroArrayTM Assay plate PM1.** Utilization of different carbon sources by *Saccharomycomorpha psychra* under controlled *in vitro* conditions, with absorbance at 590 nm measured at day 1 and day 67. Wells with a positive color change are highlighted in green.

| Well | Compound | Hit | Day | Absorption Start 590 | Absorption End 590 | Difference 590 | Mode of Action (MoA) |
|---|---|---|---|---|---|---|---|
| A01 | Negative Control | | | 0,1380 | 0,2186 | 0,0806 | C-Source, negative control |
| A02 | L-Arabinose | | 12 | 0,1142 | 0,3013 | 0,1871 | C-Source, carbohydrate |
| A03 | N-Acetyl-D-Glucosamine | | | 0,0999 | 0,1397 | 0,0398 | C-Source, carbohydrate |
| A04 | D-Saccharic Acid | | | 0,0958 | 0,1106 | 0,0148 | C-Source, carboxylic acid |
| A05 | Succinic Acid | | | 0,4060 | 0,1301 | -0,2760 | C-Source, carboxylic acid |
| A06 | D-Galactose | | | 0,1120 | 0,1482 | 0,0362 | C-Source, carbohydrate |
| A07 | L-Aspartic Acid | | | 0,0945 | 0,1324 | 0,0379 | C-Source, amino acid |
| A08 | L-Proline | | | 0,1209 | 0,1416 | 0,0207 | C-Source, amino acid |
| A09 | D-Alanine | | | 0,1269 | 0,1168 | -0,0101 | C-Source, amino acid |
| A10 | D-Trehalose | | | 0,0941 | 0,1325 | 0,0384 | C-Source, carbohydrate |
| A11 | D-Mannose | | | 0,1070 | 0,1409 | 0,0339 | C-Source, carbohydrate |
| A12 | Dulcitol | | | 0,0979 | 0,1888 | 0,0908 | C-Source, carbohydrate |
| B01 | D-Serine | | | 0,0979 | 0,1457 | 0,0478 | C-Source, amino acid |
| B02 | D-Sorbitol | | | 0,1068 | 0,1223 | 0,0155 | C-Source, carbohydrate |
| B03 | Glycerol | | | 0,0975 | 0,1226 | 0,0251 | C-Source, carbohydrate |
| B04 | L-Fucose | | | 0,0983 | 0,1527 | 0,0543 | C-Source, carbohydrate |
| B05 | D-Glucuronic Acid | | | 0,0951 | 0,1329 | 0,0378 | C-Source, carboxylic acid |
| B06 | D-Gluconic Acid | | | 0,1041 | 0,1401 | 0,0360 | C-Source, carboxylic acid |
| B07 | D,L-a-Glycerol Phosphate | | | 0,1191 | 0,1206 | 0,0015 | C-Source, carbohydrate |
| B08 | D-Xylose | | 12 | 0,1088 | 0,3401 | 0,2313 | C-Source, carbohydrate |
| B09 | L-Lactic Acid | | | 0,1131 | 0,1497 | 0,0365 | C-Source, carboxylic acid |
| B10 | Formic Acid | | | 0,0945 | 0,1363 | 0,0418 | C-Source, carboxylic acid |
| B11 | D-Mannitol | | | 0,1083 | 0,1171 | 0,0089 | C-Source, carbohydrate |
| B12 | L-Glutamic Acid | | | 0,0901 | 0,1307 | 0,0406 | C-Source, amino acid |
| C01 | D-Glucose-6-Phosphate | | | 0,0874 | 0,1152 | 0,0278 | C-Source, carbohydrate |
| C02 | D-Galactonic Acid-g-Lactone | | | 0,0907 | 0,1311 | 0,0404 | C-Source, carboxylic acid |
| C03 | D,L-Malic Acid | | | 0,1408 | 0,1382 | -0,0026 | C-Source, carboxylic acid |
| C04 | D-Ribose | | 7 | 0,1257 | 0,3961 | 0,2703 | C-Source, carbohydrate |
| C05 | Tween 20 | | | 0,2204 | 0,1351 | -0,0853 | C-Source, fatty acid |
| C06 | L-Rhamnose | | 17 | 0,1412 | 0,2162 | 0,0750 | C-Source, carbohydrate |
| C07 | D-Fructose | | | 0,0915 | 0,1169 | 0,0254 | C-Source, carbohydrate |
| C08 | Acetic Acid | | | 0,0994 | 0,1278 | 0,0284 | C-Source, carboxylic acid |
| C09 | a-D-Glucose | | | 0,0886 | 0,1150 | 0,0263 | C-Source, carbohydrate |
| C10 | Maltose | | | 0,8672 | 0,1126 | -0,7545 | C-Source, carbohydrate |
| C11 | D-Melibiose | | | 0,0970 | 0,1304 | 0,0334 | C-Source, carbohydrate |
| C12 | Thymidine | | | 0,8251 | 0,1150 | -0,7100 | C-Source, carbohydrate |
| D01 | L-Asparagine | | | 0,1043 | 0,1197 | 0,0154 | C-Source, amino acid |
| D02 | D-Aspartic Acid | | | 0,0938 | 0,1163 | 0,0226 | C-Source, amino acid |
| D03 | D-Glucosaminic Acid | | | 0,0897 | 0,1128 | 0,0231 | C-Source, carboxylic acid |
| D04 | 1,2-Propanediol | | | 0,1781 | 0,1182 | -0,0600 | C-Source, alcohol |
| D05 | Tween 40 | | | 1,2124 | 0,2166 | -0,9957 | C-Source, fatty acid |
| D06 | a-Ketoglutaric Acid | | | 0,1060 | 0,1199 | 0,0139 | C-Source, carboxylic acid |
| D07 | a-Ketobutyric Acid | | | 0,0957 | 0,1340 | 0,0383 | C-Source, carboxylic acid |
| D08 | a-Methyl-D-Galactoside | | | 0,1213 | 0,1082 | -0,0131 | C-Source, carbohydrate |
| D09 | a-D-Lactose | | | 0,1356 | 0,1165 | -0,0192 | C-Source, carbohydrate |
| D10 | Lactulose | | | 0,1120 | 0,1113 | -0,0007 | C-Source, carbohydrate |
| D11 | Sucrose | | | 0,1147 | 0,1550 | 0,0404 | C-Source, carbohydrate |
| D12 | Uridine | | | 0,0891 | 0,1696 | 0,0805 | C-Source, carbohydrate |
| E01 | L-Glutamine | | | 0,1160 | 0,1135 | -0,0025 | C-Source, amino acid |
| E02 | m-Tartaric Acid | | | 0,0894 | 0,1332 | 0,0439 | C-Source, carboxylic acid |
| E03 | D-Glucose-1-Phosphate | | | 0,0916 | 0,1159 | 0,0244 | C-Source, carbohydrate |
| E04 | D-Fructose-6-Phosphate | | | 0,1075 | 0,1392 | 0,0317 | C-Source, carbohydrate |
| E05 | Tween 80 | | | 0,6840 | 0,1161 | -0,5678 | C-Source, fatty acid |
| E06 | a-Hydroxyglutaric Acid-g-Lactone | | | 0,1099 | 0,1067 | -0,0032 | C-Source, carboxylic acid |
| E07 | a-Hydroxybutyric Acid | | | 0,0939 | 0,1048 | 0,0109 | C-Source, carboxylic acid |
| E08 | b-Methyl-D-Glucoside | | | 0,1285 | 0,1266 | -0,0018 | C-Source, carbohydrate |
| E09 | Adonitol | | | 0,1244 | 0,1103 | -0,0141 | C-Source, carbohydrate |
| E10 | Maltotriose | | | 0,0960 | 0,1273 | 0,0313 | C-Source, carbohydrate |
| E11 | 2`-Deoxyadenosine | | | 0,1186 | 0,1274 | 0,0088 | C-Source, carbohydrate |
| E12 | Adenosine | | | 0,8300 | 0,1214 | -0,7087 | C-Source, carbohydrate |
| F01 | Gly-Asp | | | 0,1033 | 0,1137 | 0,0104 | C-Source, amino acid |
| F02 | Citric Acid | | | 0,1174 | 0,1136 | -0,0039 | C-Source, carboxylic acid |
| F03 | m-Inositol | | | 0,0909 | 0,1249 | 0,0340 | C-Source, carbohydrate |
| F04 | D-Threonine | | | 0,1055 | 0,1203 | 0,0147 | C-Source, amino acid |
| F05 | Fumaric Acid | | | 0,1764 | 0,1386 | -0,0379 | C-Source, carboxylic acid |
| F06 | Bromosuccinic Acid | | | 0,0884 | 0,1156 | 0,0272 | C-Source, carboxylic acid |
| F07 | Propionic Acid | | | 0,1008 | 0,1150 | 0,0142 | C-Source, carboxylic acid |
| F08 | Mucic Acid | | | 0,4766 | 0,1130 | -0,3636 | C-Source, carboxylic acid |
| F09 | Glycolic Acid | | | 0,0934 | 0,1082 | 0,0148 | C-Source, carboxylic acid |
| F10 | Glyoxylic Acid | | | 0,0948 | 0,1230 | 0,0282 | C-Source, carboxylic acid |
| F11 | D-Cellobiose | | | 0,0903 | 0,1061 | 0,0159 | C-Source, carbohydrate |
| F12 | Inosine | | | 0,2220 | 0,1340 | -0,0880 | C-Source, carbohydrate |
| G01 | Gly-Glu | | | 0,0999 | 0,1249 | 0,0250 | C-Source, amino acid |
| G02 | Tricarballylic Acid | | | 0,0964 | 0,1295 | 0,0331 | C-Source, carboxylic acid |
| G03 | L-Serine | | | 0,1054 | 0,1259 | 0,0205 | C-Source, amino acid |
| G04 | L-Threonine | | | 0,1350 | 0,1270 | -0,0080 | C-Source, amino acid |
| G05 | L-Alanine | | | 0,1140 | 0,1372 | 0,0232 | C-Source, amino acid |
| G06 | Ala-Gly | | | 0,1135 | 0,1251 | 0,0116 | C-Source, amino acid |
| G07 | Acetoacetic Acid | | | 0,1135 | 0,1310 | 0,0174 | C-Source, carboxylic acid |
| G08 | N-Acetyl-D-Mannosamine | | | 0,1270 | 0,1193 | -0,0077 | C-Source, carbohydrate |
| G09 | Mono-Methylsuccinate | | | 0,1074 | 0,1194 | 0,0120 | C-Source, carboxylic acid |
| G10 | Methylpyruvate | | | 0,1148 | 0,1410 | 0,0262 | C-Source, ester |
| G11 | D-Malic Acid | | | 0,0875 | 0,1082 | 0,0207 | C-Source, carboxylic acid |
| G12 | L-Malic Acid | | | 0,1686 | 0,1362 | -0,0324 | C-Source, carboxylic acid |
| H01 | Gly-Pro | | | 0,1052 | 0,1517 | 0,0465 | C-Source, amino acid |
| H02 | p-Hydroxyphenyl Acetic Acid | | | 0,1411 | 0,2055 | 0,0644 | C-Source, carboxylic acid |
| H03 | m-Hydroxyphenyl Acetic Acid | | | 0,0917 | 0,1592 | 0,0675 | C-Source, carboxylic acid |
| H04 | Tyramine | | | 0,0938 | 0,1145 | 0,0207 | C-Source, amine |
| H05 | D-Psicose | | | 0,0929 | 0,1694 | 0,0765 | C-Source, carbohydrate |
| H06 | L-Lyxose | | 4 | 0,1230 | 0,4737 | 0,3507 | C-Source, carbohydrate |
| H07 | Glucuronamide | | | 0,0894 | 0,1427 | 0,0533 | C-Source, amide |
| H08 | Pyruvic Acid | | | 0,1162 | 0,1719 | 0,0557 | C-Source, carboxylic acid |
| H09 | L-Galactonic Acid-g-Lactone | | | 0,1018 | 0,1256 | 0,0238 | C-Source, carboxylic acid |
| H10 | D-Galacturonic Acid | | | 0,0919 | 0,1529 | 0,0610 | C-Source, carboxylic acid |
| H11 | Phenylethylamine | | | 0,1090 | 0,1411 | 0,0321 | C-Source, amine |
| H12 | 2-Aminoethanol | | | 0,0927 | 0,1886 | 0,0959 | C-Source, alcohol |

**Table S13 Biolog Phenotypic MicroArrayTM Assay plate PM2** Utilization of different carbon sources by Saccharomycomorpha psychra under controlled in vitro conditions, with absorbance at 590 nm measured at day 1 and day 67. Wells with a positive color change are highlighted in green.

| Well | Compound | Hit | Day | Absorption Start 590 nm | Absorption End 590 nm | Difference 590 nm | Mode of Action (MoA) |
|---|---|---|---|---|---|---|---|
| A01 | Negative Control | | | 0,1006 | 0,1549 | 0,0542 | C-Source, negative control |
| A02 | Chondroitin Sulfate C | | | 0,0914 | 0,1554 | 0,0640 | C-Source, polymer |
| A03 | a-Cyclodextrin | | | 0,0909 | 0,2034 | 0,1124 | C-Source, polymer |
| A04 | b-Cyclodextrin | | | 0,0915 | 0,1574 | 0,0659 | C-Source, polymer |
| A05 | g-Cyclodextrin | | 52 | 0,1071 | 0,1744 | 0,0673 | C-Source, polymer |
| A06 | Dextrin | | | 0,0952 | 0,1646 | 0,0694 | C-Source, polymer |
| A07 | Gelatin | | | 0,1098 | 0,1599 | 0,0501 | C-Source, polymer |
| A08 | Glycogen | | | 0,0961 | 0,1464 | 0,0503 | C-Source, polymer |
| A09 | Inulin | | | 1,1236 | 0,1888 | -0,9348 | C-Source, polymer |
| A10 | Laminarin | | | 0,0900 | 0,1832 | 0,0932 | C-Source, polymer |
| A11 | Mannan | | | 0,0909 | 0,1594 | 0,0685 | C-Source, polymer |
| A12 | Pectin | | | 0,1165 | 0,1398 | 0,0233 | C-Source, polymer |
| B01 | N-Acetyl-D-Galactosamine | | | 0,0871 | 0,1698 | 0,0827 | C-Source, carbohydrate |
| B02 | N-Acetyl-Neuraminic Acid | | | 0,1334 | 0,1576 | 0,0242 | C-Source, carboxylic acid |
| B03 | b-D-Allose | | | 0,0900 | 0,1619 | 0,0719 | C-Source, carbohydrate |
| B04 | Amygdalin | | | 0,0910 | 0,1185 | 0,0274 | C-Source, carbohydrate |
| B05 | D-Arabinose | | 11 | 0,1032 | 0,2673 | 0,1640 | C-Source, carbohydrate |
| B06 | D-Arabitol | | | 0,1687 | 0,1115 | -0,0572 | C-Source, carbohydrate |
| B07 | L-Arabitol | | | 0,1365 | 0,1193 | -0,0171 | C-Source, carbohydrate |
| B08 | Arbutin | | | 0,1084 | 0,1313 | 0,0229 | C-Source, carbohydrate |
| B09 | 2-Deoxy-D-Ribose | | 11 | 1,2684 | 0,3749 | -0,8935 | C-Source, carbohydrate |
| B10 | i-Erythritol | | | 0,0839 | 0,1295 | 0,0455 | C-Source, carbohydrate |
| B11 | D-Fucose | | | 0,0919 | 0,1303 | 0,0383 | C-Source, carbohydrate |
| B12 | 3-0-b-D-Galactopyranosyl-D-Arabinose | | 36 | 0,1155 | 0,2353 | 0,1198 | C-Source, carbohydrate |
| C01 | Gentiobiose | | | 0,0897 | 0,1472 | 0,0575 | C-Source, carbohydrate |
| C02 | L-Glucose | | | 0,0989 | 0,1324 | 0,0335 | C-Source, carbohydrate |
| C03 | D-Lactitol | | | 0,0922 | 0,1300 | 0,0378 | C-Source, carbohydrate |
| C04 | D-Melezitose | | | 0,0936 | 0,1195 | 0,0259 | C-Source, carbohydrate |
| C05 | Maltitol | | | 0,0973 | 0,1237 | 0,0264 | C-Source, carbohydrate |
| C06 | a-Methyl-D-Glucoside | | | 0,1128 | 0,1234 | 0,0106 | C-Source, carbohydrate |
| C07 | b-Methyl-D-Galactoside | | | 0,0935 | 0,1139 | 0,0204 | C-Source, carbohydrate |
| C08 | 3-Methylglucose | | | 0,0884 | 0,1514 | 0,0630 | C-Source, carbohydrate |
| C09 | b-Methyl-D-Glucuronic Acid | | | 0,9844 | 0,1172 | -0,8671 | C-Source, carboxylic acid |
| C10 | a-Methyl-D-Mannoside | | | 0,0859 | 0,1222 | 0,0363 | C-Source, carbohydrate |
| C11 | b-Methyl-D-Xyloside | | | 0,0918 | 0,1111 | 0,0193 | C-Source, carbohydrate |
| C12 | Palatinose | | 36 | 0,0973 | 0,2211 | 0,1238 | C-Source, carbohydrate |
| D01 | D-Raffinose | | | 0,1538 | 0,1361 | -0,0177 | C-Source, carbohydrate |
| D02 | Salicin | | | 0,0846 | 0,1259 | 0,0413 | C-Source, carbohydrate |
| D03 | Sedoheptulosan | | | 0,0849 | 0,1421 | 0,0573 | C-Source, carbohydrate |
| D04 | L-Sorbose | | 63 | 0,0896 | 0,1536 | 0,0640 | C-Source, carbohydrate |
| D05 | Stachyose | | | 0,0917 | 0,1381 | 0,0463 | C-Source, carbohydrate |
| D06 | D-Tagatose | | 12 | 0,0938 | 0,2014 | 0,1076 | C-Source, carbohydrate |
| D07 | Turanose | | | 0,0899 | 0,1255 | 0,0357 | C-Source, carbohydrate |
| D08 | Xylitol | | | 0,0975 | 0,1193 | 0,0218 | C-Source, carbohydrate |
| D09 | N-Acetyl-D-glucosaminitol | | | 0,7874 | 0,1137 | -0,6737 | C-Source, carbohydrate |
| D10 | g-Amino-N-Butyric Acid | | | 0,0888 | 0,1139 | 0,0251 | C-Source, carboxylic acid |
| D11 | d-Amino Valeric Acid | | | 0,0974 | 0,1041 | 0,0067 | C-Source, carboxylic acid |
| D12 | Butyric Acid | | | 0,1050 | 0,1807 | 0,0757 | C-Source, carboxylic acid |
| E01 | Capric Acid | | | 0,1092 | 0,1510 | 0,0418 | C-Source, carboxylic acid |
| E02 | Caproic Acid | | | 0,0902 | 0,1222 | 0,0320 | C-Source, carboxylic acid |
| E03 | Citraconic Acid | | | 0,1049 | 0,1107 | 0,0059 | C-Source, carboxylic acid |
| E04 | D,L-Citramalic Acid | | | 0,0925 | 0,1179 | 0,0254 | C-Source, carboxylic acid |
| E05 | D-Glucosamine | | 10 | 0,1710 | 0,3149 | 0,1438 | C-Source, carbohydrate |
| E06 | 2-Hydroxybenzoic Acid | | | 0,1549 | 0,1164 | -0,0385 | C-Source, carboxylic acid |
| E07 | 4-Hydroxybenzoic Acid | | | 0,0953 | 0,1217 | 0,0265 | C-Source, carboxylic acid |
| E08 | b-Hydroxybutyric Acid | | | 0,0943 | 0,1314 | 0,0371 | C-Source, carboxylic acid |
| E09 | g-Hydroxybutyric Acid | | | 0,1022 | 0,1293 | 0,0272 | C-Source, carboxylic acid |
| E10 | a-keto-valeric acid | | | 0,1001 | 0,1630 | 0,0629 | C-Source, carboxylic acid |
| E11 | Itaconic Acid | | | 0,1011 | 0,1049 | 0,0038 | C-Source, carboxylic acid |
| E12 | 5-Keto-D-Gluconic Acid | | 11 | 0,1146 | 0,3608 | 0,2462 | C-Source, carboxylic acid |
| F01 | D-Lactic Acid Methyl Ester | | | 0,0851 | 0,1182 | 0,0331 | C-Source, ester |
| F02 | Malonic Acid | | | 0,0858 | 0,1365 | 0,0507 | C-Source, carboxylic acid |
| F03 | Melibionic Acid | | | 0,0990 | 0,1132 | 0,0142 | C-Source, carbohydrate |
| F04 | Oxalic Acid | | | 0,1265 | 0,1138 | -0,0128 | C-Source, carboxylic acid |
| F05 | Oxalomalic Acid | | 10 | 0,1414 | 0,2946 | 0,1532 | C-Source, carboxylic acid |
| F06 | Quinic Acid | | | 0,1423 | 0,1062 | -0,0360 | C-Source, carboxylic acid |
| F07 | D-Ribono-1,4-Lactone | | | 0,1063 | 0,1243 | 0,0180 | C-Source, carboxylic acid |
| F08 | Sebacic Acid | | | 0,0855 | 0,1058 | 0,0203 | C-Source, carboxylic acid |
| F09 | Sorbic Acid | | 11 | 0,7666 | 0,3116 | -0,4550 | C-Source, carboxylic acid |
| F10 | Succinamic Acid | | | 0,0923 | 0,1126 | 0,0203 | C-Source, carboxylic acid |
| F11 | D-Tartaric Acid | | | 0,0944 | 0,1154 | 0,0210 | C-Source, carboxylic acid |
| F12 | L-Tartaric Acid | | | 0,0884 | 0,1703 | 0,0818 | C-Source, carboxylic acid |
| G01 | Acetamide | | | 0,1155 | 0,1526 | 0,0370 | C-Source, amide |
| G02 | L-Alaninamide | | | 0,0848 | 0,1153 | 0,0305 | C-Source, amide |
| G03 | N-Acetyl-L-Glutamic Acid | | | 0,0865 | 0,1121 | 0,0256 | C-Source, amino acid |
| G04 | L-Arginine | | | 0,0892 | 0,1288 | 0,0396 | C-Source, amino acid |
| G05 | Glycine | | | 0,1585 | 0,1498 | -0,0087 | C-Source, amino acid |
| G06 | L-Histidine | | | 0,1400 | 0,1360 | -0,0040 | C-Source, amino acid |
| G07 | L-Homoserine | | | 0,0962 | 0,1194 | 0,0232 | C-Source, amino acid |
| G08 | Hydroxy-L-Proline | | | 0,0973 | 0,1215 | 0,0242 | C-Source, amino acid |
| G09 | L-Isoleucine | | | 0,6810 | 0,1272 | -0,5538 | C-Source, amino acid |
| G10 | L-Leucine | | | 0,0947 | 0,1317 | 0,0370 | C-Source, amino acid |
| G11 | L-Lysine | | | 0,1013 | 0,1207 | 0,0194 | C-Source, amino acid |
| G12 | L-Methionine | | | 0,1293 | 0,1747 | 0,0454 | C-Source, amino acid |
| H01 | L-Ornithine | | | 0,0996 | 0,1601 | 0,0605 | C-Source, amino acid |
| H02 | L-Phenylalanine | | | 0,0950 | 0,1699 | 0,0749 | C-Source, amino acid |
| H03 | L-Pyroglutamic Acid | | | 0,1701 | 0,1542 | -0,0159 | C-Source, amino acid |
| H04 | L-Valine | | | 0,0861 | 0,1405 | 0,0544 | C-Source, amino acid |
| H05 | D,L-Carnitine | | | 0,0999 | 0,1263 | 0,0264 | C-Source, carboxylic acid |
| H06 | Sec-Butylamine | | | 0,0950 | 0,1394 | 0,0444 | C-Source, amine |
| H07 | D,L-Octopamine | | | 0,0879 | 0,1289 | 0,0409 | C-Source, amine |
| H08 | Putrescine | | | 0,1840 | 0,1623 | -0,0217 | C-Source, amine |
| H09 | Dihydroxyacetone | | 4 | 0,8661 | 0,5639 | -0,3023 | C-Source, alcohol |
| H10 | 2,3-Butanediol | | | 0,0888 | 0,1328 | 0,0440 | C-Source, alcohol |
| H11 | 2,3-Butanone | | | 0,0892 | 0,1563 | 0,0672 | C-Source, alcohol |
| H12 | 3-Hydroxy 2-Butanone | | | 0,0996 | 0,1768 | 0,0772 | C-Source, alcohol |

**Data S1. High-confidence carbohydrate-active enzymes (CAZymes) predicted in the *Saccharomycomorpha psychra* genome.** The table includes gene identifiers, CAZyme family assignments inferred by HMMER, DIAMOND and Hotpep (dbCAN2), SignalP secretion predictions, and the number of supporting annotation tools used for confidence assessment.

**Data S2. High-confidence membrane transporter annotation of *Saccharomycomorpha psychra*.** The table contains gene identifiers, best TCDB assignments, UniProt homology, PFAM domain annotations, predicted transmembrane helices, annotation confidence, transporter class and subclass, transport mechanism, and transporter family classification.

**Data S3. Predicted secretome of *Saccharomycomorpha psychra*.** The table includes gene identifiers, signal peptide predictions (SignalP and Phobius), transmembrane topology, predicted subcellular localization, GPI-anchor predictions, confidence scores, and the final classification of secreted proteins.
